# Structure-based analysis of CysZ-mediated cellular uptake of sulfate

**DOI:** 10.1101/131318

**Authors:** Zahra Assur Sanghai, Qun Liu, Oliver B. Clarke, Edgar Leal Pinto, Brian Kloss, Shantelle Tabuso, James Love, Marco Punta, Surajit Banerjee, Kanagalaghatta R. Rajashankar, Burkhard Rost, Diomedes Logothetis, Matthias Quick, Wayne A. Hendrickson, Filippo Mancia

## Abstract

Sulfur, most abundantly found in the environment as sulfate (SO_4_^2-^), is an essential element in metabolites required by all living cells, including amino acids, co-factors and vitamins. Current understanding of the cellular delivery of SO_4_^2-^ at the molecular level is limited however. CysZ has been described as a SO_4_^2-^ permease, but its sequence family is without known structural precedent. Based on crystallographic structure information, SO_4_^2-^ binding and uptake experiments in cells and proteoliposomes, and single-channel conductance measurements, we provide insight into the molecular mechanism of CysZ-mediated translocation of SO_4_^2-^ across membranes. CysZ properties differ markedly from those of known transporters and ion channels. The structures display a hitherto unknown fold with dual topology, assembling in CysZ from *Pseudomonas denitrificans* as a trimer of antiparallel dimers in the membrane. CysZ structures from two other species recapitulate dimers from this assembly. Mutational studies highlight the functional relevance of conserved CysZ residues.

---

Sulfur has a central role in many cellular processes across all kingdoms of life. It is a vital component of several essential compounds, including the sulfur-containing amino acids cysteine and methionine, in prosthetic groups such as the Fe-S clusters, as well as vitamins and micronutrients such as biotin (vitamin H), thiamine (vitamin B1) and lipoic acid, and in coenzymes A and M (Barton, 2005). Whilst mammals obtain the majority of the necessary sulfur-containing metabolites directly from the diet, plants, fungi, and bacteria are able to assimilate and utilize sulfur from organic and inorganic sources (Barton, 2005). Sulfate (SO_4_^2-^) is the most abundant source of sulfur in the environment and its utilization is contingent upon its entry into the cell (Kertesz, 2000). In certain fungi, and prokaryotes, once internalized, SO_4_^2-^ is first reduced to sulfite (SO_3_ ^2-^), and then further to sulfide (S^2-^), a form that can be used by the cell (Kredich, Hulanicka, & Hallquist, 1979) (Fig. 1-S1). In *Escherichia coli* and other gramnegative bacteria, the culmination of the aforementioned sulfate assimilatory (also known as reductive) pathway is the formation of cysteine by the addition of S^2-^ to O-acetylserine by cysteine synthase, followed by the synthesis of methionine from homocysteine (Kredich, 1971) (Fig. 1-S1).

**Fig. 1.**
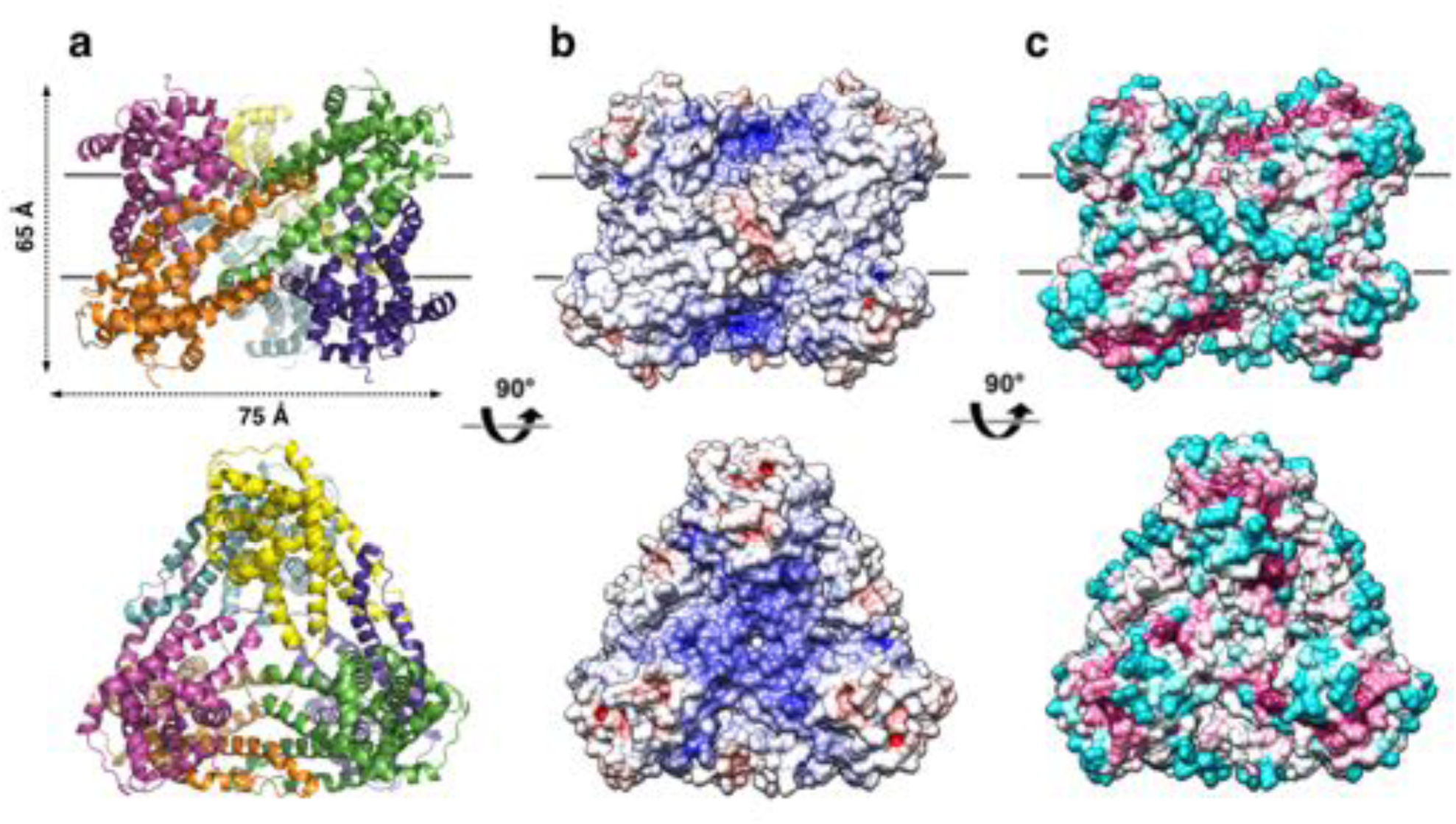
Overall structure of the *P. denitrificans* CysZ (*Pd*CysZ) hexamer. a. Side and top views of the hexamer as a ribbon diagram with each protomer chain colored differently. The approximate dimensions of the hexamer marked in Å. b. Side and top views represented by surface electrostatics as calculated by APBS, with negative and positive surface potential represented in red and blue respectively. c. Side and top views representing conservation of residues as calculated by ConSurf, with maroon being most conserved to cyan being least conserved.

In prokaryotes, the entry of SO_4_^2-^ into the cell is mediated by four known families of dedicated transport systems: the ABC sulfate transporter complexes SulT or CysTWA, the SulP family of putative SLC13 sodium:sulfate or proton:sulfate symporters or SLC26 solute:sulfate exchangers, the phosphate transporter-like CysP/PitA family, and the CysZ family classified as SO_4_^2-^ permeases (Aguilar-Barajas, Diaz-Perez, Ramirez-Diaz, Riveros-Rosas, & Cervantes, 2011; Hryniewicz, Sirko, Palucha, Bock, & Hulanicka, 1990; Kertesz, 2001; Loughlin, Shelden, Tierney, & Howitt, 2002; Mansilla & de Mendoza, 2000; Sirko, Zatyka, Sadowy, & Hulanicka, 1995). CysZ family members are 28-30 kDa bacterial inner-membrane proteins found exclusively in prokaryotes with no apparent homology to any of the established channel or transporter folds, and are scarcely studied in the literature (Zhang, Jiang, Nan, Almqvist, & Huang, 2014). The *cysZ* gene owes its name to its presence in the cysteine biosynthesis regulon. In two reports from thirty years ago, an *E. coli* K12 strain with a *cysZ* deletion showed a severe impairment in its ability to accumulate SO_4_^2-^ and was not viable in sulfate-free media without an alternate sulfur source such as thiosulfate (S_2_O_3_ ^2-^) (Britton et al., 1983; Parra, Britton, Castle, Jones-Mortimer, & Kornberg, 1983). Recently, a third report studying the functional properties of CysZ, concluded that the protein from *E. coli* functions as a high affinity, highly specific pH-dependent SO_4_^2-^ transporter, directly regulated by the toxic, assimilatory pathway intermediate, SO_3_ ^2-^ (Zhang et al., 2014).

To investigate the role of CysZ in cellular sulfate uptake at a molecular level, we have undertaken an approach that combines structural and functional studies. To this end, we determined the crystal structures of CysZ from three species, *Idiomarina loihiensis (Il; IlCysZ)*, *Pseudomonas fragi (Pf; PfCysZ),* and *Pseudomonas denitrificans (Pd; PdCysZ)*, and characterized CysZ function in purified form, in reconstituted proteoliposomes, in planar lipid bilayers, and in cells. Combining the structural information from the three orthologs reveals that CysZ features a novel protein fold that assembles as oligomers with a dual topology. This arrangement can be understood as being derived from trimers of dimers akin to the hexameric assembly captured in one of the structures. Interpreting the functional data in a structural context has allowed us to formulate a mechanistic model for CysZ-mediated SO_4_^2-^ translocation across the bacterial cytoplasm membrane.

Both the structures and the functional properties of CysZ proteins are distinct from those of any known membrane transporter or ion channel. Besides not resembling other transporter structures, CysZ mediates sulfate uptake into cells or proteoliposomes without coupling to ion gradients, partner proteins, or exogenous energy sources such as ATP. Besides differing from other ion channel structures, CysZ generates sulfate currents across lipid bilayers with unusual characteristics. These distinctive properties make CysZ appealing as a model system for studies of biophysical principles of membrane protein biogenesis and transmembrane ion passage.

## RESULTS

### Structure determination of CysZ

Following a structural genomics approach aimed at crystallization for structural analysis, we cloned and screened a total of 63 different bacterial homologs of CysZ for high-level expression and stability in detergents (Love et al., 2010; Mancia & Love, 2011). Crystal structures were determined for CysZ from three organisms. Chronologically, the structure of *Il*CysZ was the first solved, to 2.3 Å resolution in space group *C2* by SAD, initially based on a single selenate ion bound to the protein and subsequently also by selenomethione derivatization (SeMet) SAD, and multi-crystal native SAD (Liu et al., 2012). The structure of *Pf*CysZ was solved second, to 3.5 Å resolution, with crystals also belonging to space group *C2.* Although *Pf*CysZ and *Il*CysZ share 42% sequence identity, molecular replacement failed to find a convincing solution, and we instead used SeMet-derivatized *Pf*CysZ to obtain phase information by multi-crystal SeMet SAD. Third, we determined the structure of *Pd*CysZ, which crystallized in multiple forms belonging to space groups *P6*_*3*_, *P4*_*1*_*22* and *P2*_*1*_*2*_*1*_*2*_*1*_, revealing the same architecture and oligomeric assembly each time (Fig. 1-S2b). *Pd*CysZ structures in the *P6*_*3*_ and *P2*_*1*_*2*_*1*_*2*_*1*_ lattices each contain an entire hexamer in their asymmetric units, whereas a molecular diad coincides with a crystallographic axis in the *P4*_*1*_*22* lattice. We focused our analysis on the best of these (3.4 Å resolution in *P6*_*3*_). The location of the selenium sites was obtained from a SeMet SAD data set, and the structure was solved by combining the resulting SAD phases with those from a molecular replacement solution obtained by positioning the *Pf*CysZ model (75% sequence identity) onto SeMet fiducials in the initial electron density map (Table 1, Fig. 1-S2a).

**Table 1:** Crystallographic data and refinement statistics. Values in parentheses are from the highest resolution shell. R_free_ was calculated using 5% of data excluded from refinement.

| Data collection | Native *IlCysZ* | *IlCysZ*<br>w/$\text{SeO}_4^{2-}$<br>(SAD) | SeMet *PfCysZ*<br>(6 crystals,<br>refinement) | SeMet<br>*PfCysZ*<br>(15 crystals,<br>SAD) | Native<br>*PdCysZ*<br>(MR-SAD) |
| --- | --- | --- | --- | --- | --- |
| **Beamline** | NSLS X4A | NSLS X4A/C | APS 24-ID-E | APS 24-ID-C | APS 24-ID-C |
| **Space group** | C2 | C2 | C2 | C2 | P6<sub>3</sub> |
| **Cell dimensions:** |  |  |  |  |  |
| **a, b, c (Å)** | 128.9, 82.0,<br>100.4 | 128.9, 81.9, 100.3 | 172.29, 56.9, 96.17 | 172.35, 56.9, 96.31 | 225.13, 225.13,<br>96.62 |
| **$\alpha, \beta, \gamma$ (°)** | 90, 125.1, 90 | 90, 125.1, 90 | 90, 91.43, 90 | 90, 91.33, 90 | 90, 90, 120 |
| **Z<sub>a</sub>** | 2 | 2 | 2 | 2 | 6 |
| **Wavelength** | 1.7432 | 0.96789 | 0.97890 | 0.97890 | 1.0230 |
| **Bragg spacings (Å)** | 30-2.30 | 50-2.10 | 40-3.20 | 40-3.50 | 86-3.40 |
| **R<sub>merge</sub>** | 0.047 (0.257) | 0.046 (0.417) | 0.136 (6.506) | 0.135 (2.13) | 0.095 (2.89) |
| **I / σ<sub>I</sub>** | 29.4 (7.9) | 25.0 (1.9) | 12.3 (0.7) | 24.2 (2.6) | 22.4 (1.2) |
| **Completeness (%)** | 99.9 (99.9) | 99.9 (100.0) | 99.8 (99.3) | 99.9 (100.0) | 100.0 (100.0) |
| **Multiplicity** | 7.3 (7.2) | 7.6 (7.6) | 27.2 | 72.8 | 19.7 |
| **Refinement:** |  |  |  |  |  |
| **Resolution (Å)** | 2.30 |  | 3.50 |  | 3.40 |
| **No. of reflections** | 38075 |  | 20741 |  | 37221 |
| **R<sub>work</sub> / R<sub>free</sub>** | 0.200/0.238 |  | 0.297/0.339 |  | 0.239/0.277 |
| **No. of atoms:** |  |  |  |  |  |
| **Protein** | 3679 |  | 3428 |  | 11415 |
| **Ligand/ion** | 249 |  | 0 |  | 560 |
| **Water** | 162 |  | 0 |  | 0 |
| **Average B-factors (Å<sup>2</sup>)** | 49.4 |  | 198.9 |  | 204.18 |
| **Protein** | 48.4 |  | 198.9 |  | 204.18 |
| **Ligand/Ion** | 77.3 |  | - |  | - |
| **Water** | 45.3 |  | - |  | - |
| **Bond Ideality (r.m.s.d.):** |  |  |  |  |  |
| **Bond lengths (Å)** | 0.006 |  | 0.006 |  | 0.004 |
| **Bond angles (°)** | 0.894 |  | 1.133 |  | 0.96 |
| **Ramachandran** |  |  |  |  |  |
| **Analysis:** |  |  |  |  |  |
| **Favored (%)** | 99.0 |  | 99.2 |  | 98.5 |
| **PDB accession code** | 3TX3 |  | TBD |  | TBD |

### The hexameric structure of *Pd*CysZ

The refined structure of *Pd*CysZ comprises an entire hexamer of near-perfect D3 symmetry. Antiparallel pairs of protomers arrange together as a trimer of dimers (Fig. 1a), with the three-fold axis oriented perpendicular to the plane of the putative membrane and three two-fold axes between dimers of the hexamer. Both the periplasmic and cytoplasmic faces of the hexamer are essentially identical by symmetry, resulting in a dual-topology assembly for *Pd*CysZ. The hexamer has a triangular face of equal sides measuring approximately 75 Å, with the perpendicular span of about 65 Å. The interaction of the six protomers results in a total buried surface area (Krissinel & Henrick, 2007) of 5,700 Å^2^. A surface electrostatic representation reveals a hydrophobic belt along the mid-section of the hexamer when viewed from its side, outlining the orientation of CysZ in the lipid bilayer (Czodrowski, Dramburg, Sotriffer, & Klebe, 2006; Dolinsky et al., 2007; Dolinsky, Nielsen, McCammon, & Baker, 2004) (Fig. 1b). A surface representation of the sequence conservation, calculated by analysis of multiple sequence alignments (MSA) (Ashkenazy et al., 2016; Glaser et al., 2003) highlights the regions of invariance in the sequence and in turn, the areas on the molecule that are most likely to have structural and functional importance (Fig. 1c).

The CysZ protomer is an alpha-helical integral membrane protein with two long transmembrane (TM) helices (H2b and H3a) and two pairs of shorter helices (H4b-H5a and H7-H8) that insert only partially into the membrane (hemi-penetrating), forming a funnel or tripod-like shape within the membrane (Fig. 2a, b). The protein has an extra-membranous hydrophilic ‘head’, comprising an iris-like arrangement of the two short helices, H1 and H6, and kinked helices H3b, H4a, and H5b. The amino and carboxyl termini are also located in this region (Fig. 2a, b). H4b and H5a lie partly inserted in the membrane, with the turn between the two helices pointing in towards the three-fold axis at the center of the hexamer. CysZ amino-acid sequences from different organisms are very similar; for example, those of *Pd*CysZ and *E. coli* CysZ (*Ec*CysZ) are 40.5% identical and those of *Ec*CysZ and our three structures have 30.0% of their residues exactly in common (Fig. 2c). Helix boundaries are also essentially the same in the structures of *Pd*CysZ, *Il*CysZ and *Pf*CysZ (Fig. 2c).

**Fig. 2.**
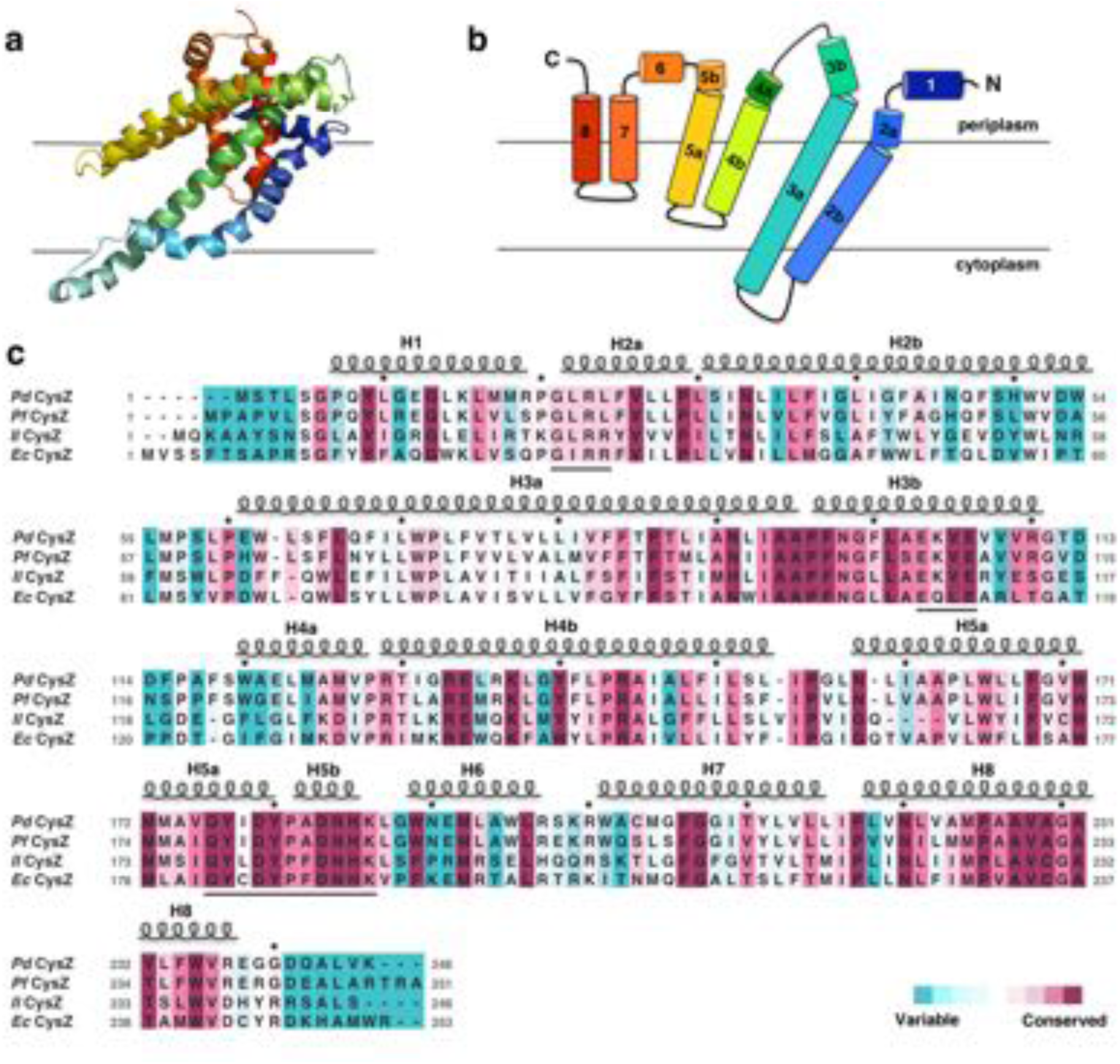
Structure and topology diagram of *Pd*CysZ protomer, with sequence alignment and conservation. a. Ribbon diagram of the *Pd*CysZ protomer colored in rainbow colors from N (blue) to C terminus (red), viewed from within the plane of the membrane, shown in the same orientation as the protomer drawn in green in Fig. 1a. b. Topology diagram of the *Pd*CysZ protomer with helices marked from 1 to 8. Helices H2b and H3a are transmembrane helices, whereas H4-H5 and H7-H8 are hemi-penetrating helical hairpins, only partially inserted into the membrane. c. Sequence alignment of *E. coli* CysZ (*Ec*CysZ), *P. denitrificans* CysZ (*Pd*CysZ), *P. fragi* CysZ (*Pf*CysZ), and *I. loihiensis* CysZ (*Il*CysZ). Residues are colored based on conservation, with maroon being most conserved and cyan least conserved, as calculated by ConSurf using a sequence alignment of 150 non-redundant sequences from the CysZ family as input. Spirals above residues mark the extent of the helical segments based on the atomic structure of *Pd*CysZ with helices numbered H1-H8; letters mark residue identities; black dots above identify every tenth residue (modulo 10) in the *Pd*CysZ sequence and black underlines mark functionally relevant motifs discussed in the text.

### The dimeric assembly of *Il*CysZ and *Pf*CysZ

Unlike *Pd*CysZ, both *Il*CysZ and *Pf*CysZ crystallize as dimers (Fig. 3a, b), in agreement with their different behavior in detergent-containing solution. Indeed, size-exclusion chromatography runs of the three species of CysZ in the same buffer and detergent conditions, show mono-disperse peaks eluting at 13.42 ml (*Il*CysZ), 13.95 ml (*Pf*CysZ) and 12.56 ml (*Pd*CysZ) (Fig. 3-S1). This result is consistent with a different and smaller oligomeric state of *Il*CysZ and *Pf*CysZ (similar retention volume) compared to *Pd*CysZ (eluting 1 ml ahead).

**Fig. 3.**
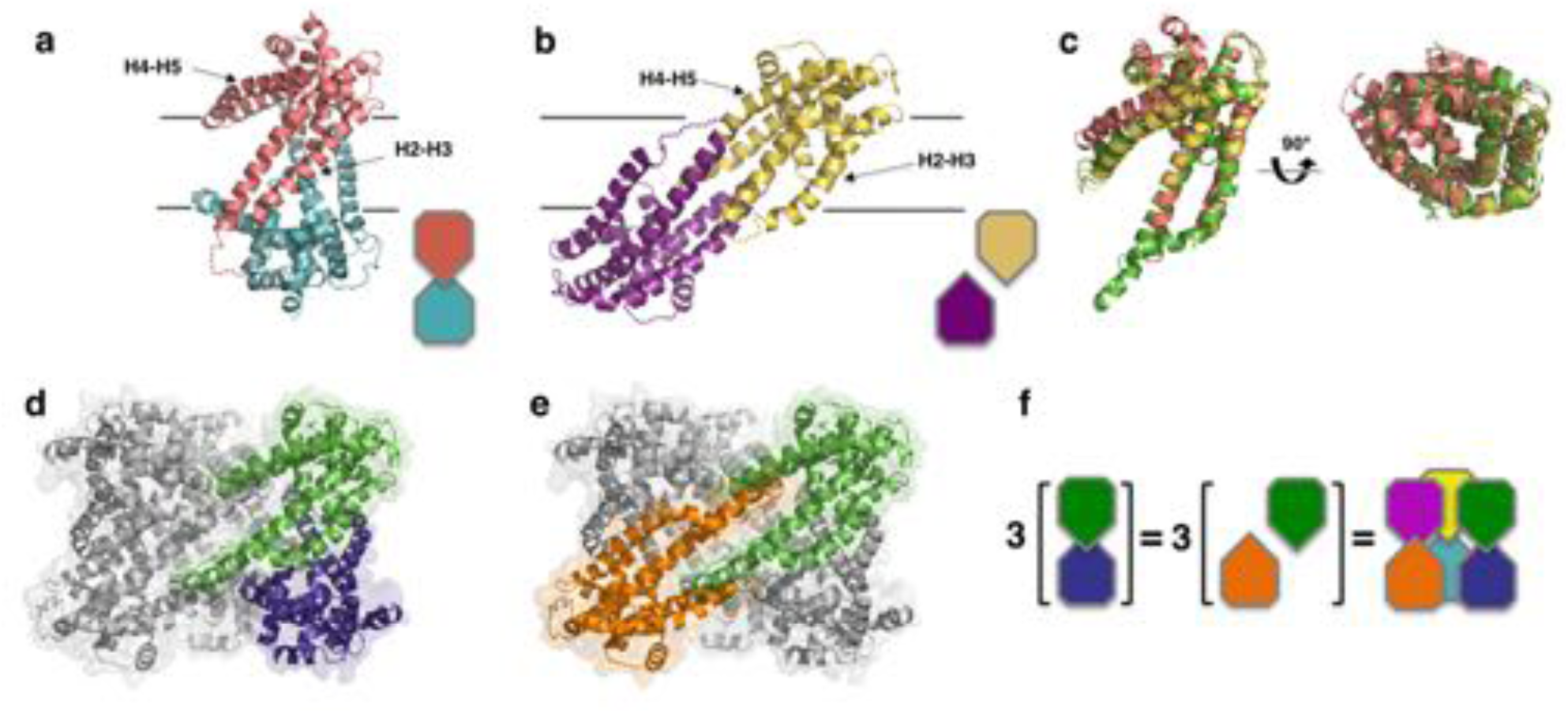
Structures of *Il*CysZ and *Pf*CysZ, with comparison to *Pd*CysZ. a. Ribbon diagram of the structure of CysZ from *I. loihiensis (Il*CysZ) at 2.3 Å. Protomers of the dimer are colored in salmon pink and teal blue, arranged in a head-to-tail conformation in the membrane, with helical hairpins H2-3 and H4-5 labeled for clarity. The dimer interface of *Il*CysZ involves H2-3. b. Ribbon diagram of the structure of CysZ from *P. fragi* (*PfCysZ*) at 3.2 Å. The protomers of the dual topology dimer in the membrane are colored gold and purple, with helical hairpins H2-3 and H4-5 again labeled. The dimer interface here involves the interaction of helices H4-5 of each protomer. c. Side and top views of the superposition of the three different protomers from *Pd*CysZ (green), *Il*CysZ (pink) and *Pf*CysZ (yellow) after aligning H1-H3. d, e. The same dimer interfaces observed in *Il*CysZ (a) and *Pf*CysZ (b) observed in the hexameric assembly of *Pd*CysZ, as highlighted in green and blue (d) and green and orange (e). f. Schematic representation showing how three copies of the dimeric protomers of *Il*CysZ (green and blue, left) and of *Pf*CysZ (orange and green, center), can coexist in and each recapitulate the hexameric assembly of *PdCysZ*.

The protomers of the *Il*CysZ dimer are arranged in a head-to-tail antiparallel association, with H4b-H5a protruding at an angle that is nearly parallel to the putative plane of the membrane (Fig. 3a, 3-S2a, b). The *Pf*CysZ dimer is also arranged in an antiparallel orientation but with a different dimer interface (Fig. 3b). In the *Pf*CysZ structure, H4b-H5a together form a narrower angle with H2 and H3, and tuck-in closer to the rest of the molecule (Fig. 3b). The resulting dumbbell-shaped *Pf*CysZ dimer is predicted, by OPM/PPM (Orientation of Proteins in Membranes) (Lomize, Pogozheva, Joo, Mosberg, & Lomize, 2012), to lie in the membrane at a 31° tilt to the perpendicular, in agreement with the position of its central hydrophobic belt, as revealed by surface-electrostatics calculations (Fig. 3-S2e, f). The dimer interfaces of *IlCysZ* and *PfCysZ* bury 780 Å^2^ and 1136 Å^2^ of surface area respectively (Krissinel & Henrick, 2007)

The individual protomers of CysZ from all three species adopt the same topology and fold, and superpose well with an overall pairwise root mean squared deviation (r.m.s.d) of ∼2.5 Å (Fig. 3c). The greatest variation between the protomers of each structure is seen in the orientation of H4b-H5a with respect to the TM helices, H2 and H3 (Fig. 3c), which seem to be the most conformationally flexible with respect to the rest of the molecule.

Comparison of the structures of *Il*CysZ and *Pf*CysZ with that of *Pd*CysZ revealed that these two distinctive dimeric structures are both represented in the hexameric one. Indeed, the *Il*CysZ dimer (Fig. 3a) resembles the vertically arranged pair of protomers in the *Pd*CysZ hexamer (Fig. 3d) at each of the vertices of the triangular structure. On the other hand, the *Pf*CysZ dimer (Fig. 3b) resembles the transverse pair of protomers lying at a tilt, just as was predicted by OPM (Lomize et al., 2012), along the side of the *Pd*CysZ hexamer when viewed from inside the plane of the membrane (Fig. 3e). In essence, the *Pd*CysZ hexamer can be seen as a trimer of either *Il*CysZ or *Pf*CysZ dimers (Fig. 3f).

### Dual-topology assembly of CysZ

To validate the dual-topology assembly of CysZ observed in all our structures, we performed disulfide crosslinking assays on engineered cysteine mutants of CysZ designed to capture the antiparallel dimer, utilizing isolated membranes. Disulfide-trapping experiments of the transverse dimer, performed by crosslinking a pair of mutants (L161C-A164C) on H5 of *Pf*CysZ, as well as the corresponding pair in *Il*CysZ (V157C-Q163C) confirm the dimer interface observed in the structures of *Pf*CysZ and *Il*CysZ, and, as a consequence, the dual-topology assembly of the two proteins (Fig. 3-S2f, g, h).

To further validate the dual-topology orientation of CysZ in the membrane, we performed a cysteine accessibility scan experiment, mapping residues expected to be located outside the lipid bilayer, by fluorescence labeling of the thiol groups of various single cysteine mutants with a membrane-impermeable dye. Membrane fractions isolated from recombinant cultures expressing *Il*CysZ cysteine mutants introduced at positions along the edge of helix H4 predicted to be solvent accessible based on our structure, were indeed labeled with a membrane-impermeable fluorescent thiol-specific maleimide dye (Fig. 3-S2c, d).

### Functional characterization of CysZ

To characterize the functional properties of CysZ, we used a four-pronged approach: (i) radiolabeled [^35^S]O_4_^2-^ uptake experiments in cells expressing plasmid-encoded CysZ, (ii) radiolabeled ([^35^S]O_4_^2-^) binding to purified CysZ in detergent solution, (iii) [^35^S]O_4_^2-^ uptake in proteoliposomes reconstituted with purified CysZ, and (iv) single-channel electrophysiological recordings in a planar lipid bilayer reconstituted with CysZ.

First, we compared the time course of SO_4_^2-^ accumulation in an *E. coli cysZ* knockout strain (*E. coli* K12 JW2406-1, CysZ^*-*^) (Baba et al., 2006) with that in the wild-type (WT, *E. coli* K12 BW25113, *cysZ*^*+*^) strain. CysZ^-^ cells, after growth in minimal media and 12 hours of sulfate starvation, showed significantly diminished SO_4_^2-^ uptake when compared to the WT strain, consistent with previous results (Fig. 4a) (Parra et al., 1983). The CysZ^-^ strain still showed a low level of sulfate accumulation, which could be attributed to the other endogenous sulfate transport systems present in the bacteria (for example ABC transporter and SulP). This uptake-deficient phenotype of the CysZ^-^ cells could be rescued by plasmid-driven expression of *Pd*CysZ (Fig. 4b), as well as *Il*CysZ and *Pf*CysZ (data not shown). Consistent with previously reported data (Zhang et al., 2014), we observed that SO_3_ ^2-^ ions severely impede or block the accumulation of [^35^S]O_4_^2-^ by CysZ-expressing cells (Fig. 4b).

**Fig. 4.**
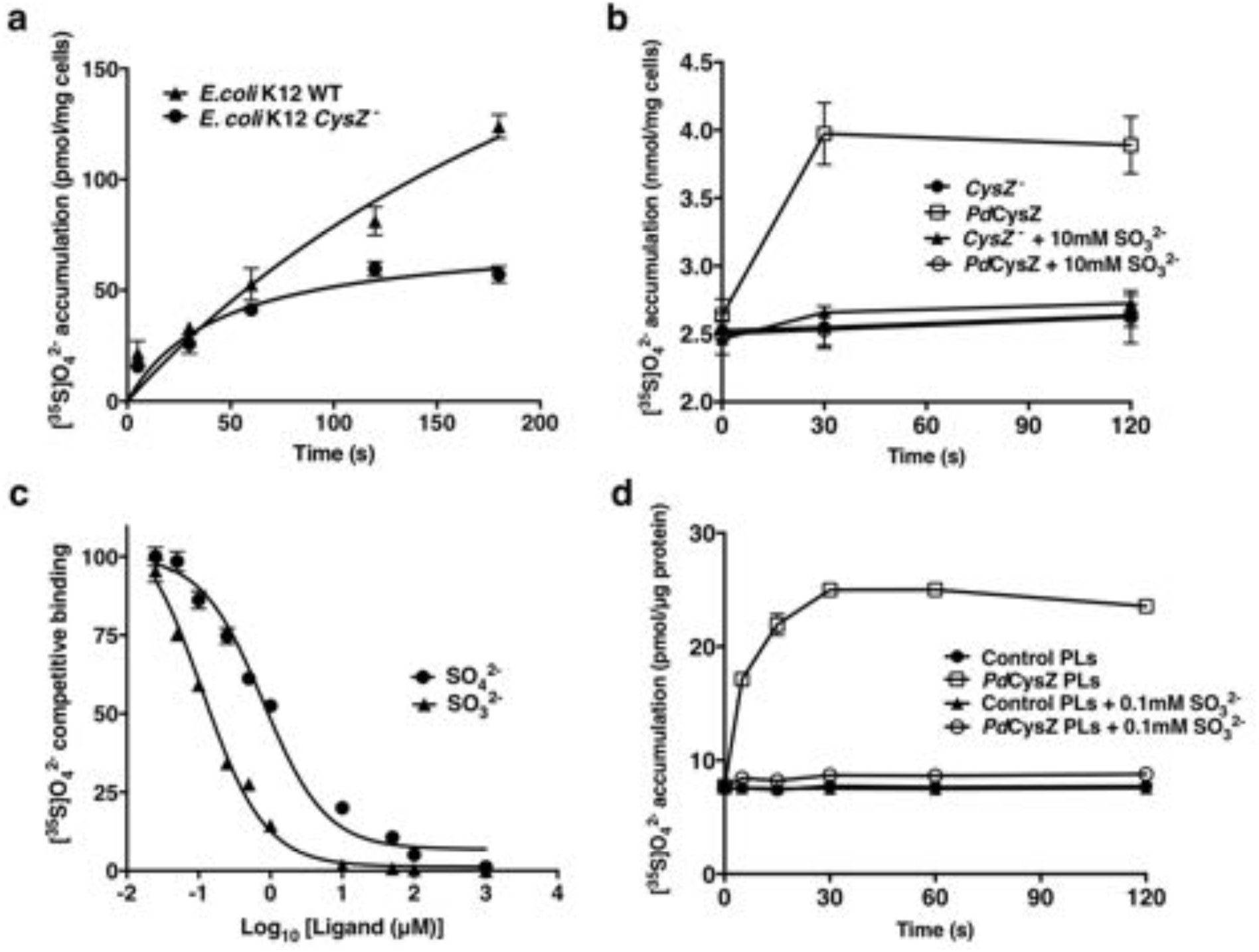
Functional characterization of CysZ. a. Time course of [^35^S]O_4_^2-^ uptake (320 μM) by cells of WT *E. coli K12* (strain BW25113) and by CysZ^-^ (strain JW2406-1) cells (n=3). b. Sulfite inhibition of sulfate uptake. Radiolabeled sulfate (320 μM [^35^S]O_4_^2-^) uptake is rescued in the same *cysZ*^*-*^ knockout strain by transformation with an expression vector for *Pd*CysZ whereas the control cells were transformed with an empty vector. Sulfate uptake is inhibited by the addition of 10 mM SO_3_ ^2-^ (n=3). c. Competitive binding of labeled sulfate in presence of unlabeled ligands. The inhibition of labeled sulfate (7 nM [^35^S]O_4_^2-^) binding by non-labeled sulfate or sulfite is measured by the scintillation proximity assay using purified *Pd*CysZ (n=3). *Pd*CysZ binds SO_4_^2-^ with an *EC*_*50*_ of 0.81 ± 0.045 μM and exhibits an *IC*_*50*_ of 0.12 ± 0.034 μM for SO_3_ ^-2^. d. Radiolabeled sulfate (10 μM [^35^S]O_4_^2-^) accumulation into proteoliposomes reconstituted with detergent-purified *Pd*CysZ. Uptake into reconstituted proteoliposomes (open squares) is shown in comparison with control (empty) liposomes (solid discs). Uptake is inhibited by the presence of 0.1 mM SO_3_ ^2-^ (n=3).

To further characterize the interaction of CysZ with both SO_4_^2-^ and SO_3_ ^2-^, we performed binding experiments on purified *Pd*CysZ using the scintillation proximity assay (SPA) (Quick & Javitch, 2007). To determine the concentration of half-maximal binding (*EC*_*50*_), we isotopically diluted [^35^S]O_4_^2-^ with non-labeled SO_4_^2-^, obtaining an *EC*_*50*_ of 0.81 ± 0.045 μM (Fig. 4c). Competing binding of [^35^S]O_4_^2-^ with SO_3_ ^2-^ revealed that *Pd*CysZ binds SO_3_ ^2-^ with greater affinity as reflected by a half-maximum inhibition constant (*IC*_*50*_) of 0.12 ± 0.034 μM.

Next, we conducted [^35^S]O_4_^2-^ uptake experiments on detergent solubilized and purified CysZ reconstituted in proteoliposomes. Proteoliposomes provide a means for assessing the activity of a membrane protein independent of other proteins found natively in cellular expression systems. Such binding partners could be involved in SO_4_^2-^ uptake as measured in intact *E. coli* cells (Fig. 4a). Furthermore, in addition to the SPA measurements, these experiments allowed us to confirm that the detergent-based extraction from the membrane and purification of CysZ used for our crystallization approaches did not compromise its activity. Acting alone, *Pd*CysZ in proteoliposomes mediated the accumulation of SO_4_^2-^ in a time-dependent manner; the uptake occurred rapidly, peaking at about 30 seconds after incubation with the radioligand (Fig. 4d). Consistent with our observations on cells, the influx of sulfate was inhibited by the presence of SO_3_ ^2-^ at micromolar concentrations. *Il*CysZ and *Pf*CysZ exhibited similar uptake profiles in liposomes, suggesting similar functional properties (Fig. 4-S1a).

To determine whether sulfate uptake by CysZ was driven by the concentration gradient of sulfate itself, or if it was coupled to a secondary driving force such as an ion (e.g., H^+^, Na^+^) gradient, we tested the effect of the uncouplers CCCP (carbonyl cyanide m-chlorophenyl hydrazone), and oligomycin, as well as that of the ionophores valinomycin and gramicidin on CysZ-mediated [^35^S]O_4_^2-^ accumulation in our cell-based assay (Buckler & Vaughan-Jones, 1998). Since none of these uncoupling agents had a significant effect on the accumulation of [^35^S]O_4_^2-^, these experiments suggest that CysZ-mediated [^35^S]O_4_^2-^ flux is not dependent on the ‘classical’ H^+^ or Na^+^ electrochemical transmembrane gradient, thus indicating that the observed uptake solely depends on the SO_4_^2-^ transmembrane concentration gradient (Fig. 4-S1b), a notion that is consistent with a channel-like mechanism for CysZ-mediated SO_4_^2-^ translocation.

Finally, we employed the use of single-channel electrophysiological current recordings on CysZ (*Pd*CysZ and *Il*CysZ) reconstituted in a planar lipid bilayer (Fig. 4-S2). Consistent with our [^35^S]O_4_^2-^-based flux assays, these electrophysiological measurements showed that reconstituted CysZ exhibits channel-like properties when a membrane potential is applied in symmetrical Na_2_SO_4_ solutions on both sides of the bilayer. The *Il*CysZ protein exhibits an average open probability (P_OPEN_) of 0.8 and conductance estimated as 92 ± 12 pS. However, unitary conductance levels varied, showing a dependence on the amount of protein reconstituted in the planar lipid bilayer. The *Pd*CysZ protein passes a broad distribution of currents at fixed voltages. For both proteins, the SO_4_^2-^ currents are completely abolished by the presence of low micromolar SO_3_ ^2-^ concentrations.

### Sulfate Binding Site

A bound SO_4_^2-^ ion was observed in the structure of *Il*CysZ. This binding site was confirmed by purifying and crystallizing CysZ in the presence of the heavier SO_4_^2-^-analog selenate (SeO_4_^2-^). In the SO_4_^2-^ bound structure, each protomer in the dimer showed electron density consistent with a bound sulfate, but only one of the two refined to full occupancy, hence only one is shown in the refined model of *Il*CysZ (Fig. 5a). The location of the bound SO_4_^2-^ is close to the putative membrane interface where it is coordinated by two arginine residues (R27 and R28 in *Il*CysZ) and the backbone amides of residues G25 and L26. This SO_4_^2-^-binding site is in the loop between H1 and H2, with the motif GLR(R) being well conserved among the CysZ family. Consistent with this observation, crystals of *Il*CysZ in which R27 and R28 were replaced with alanines did not show interaction with SO_4_^2-^ in that site based on i) the lack of any density in the refined mutant structure at 2.3 Å and ii) functional studies (see below). We could not confirm the presence of SO_4_^2-^ in either of the other structures of *Pf*CysZ and *Pd*CysZ, owing, at least in part, to their lower resolution limit (∼3.5 Å). Soaking and co-crystallization attempts on both *Pf*CysZ and *Pd*CysZ with SeO_4_^2-^ resulted in cracking and destabilization of crystals, and loss of measurable diffraction. In this vein, *Il*CysZ R27A/R28A, and the corresponding *Pf*CysZ R25A and *Pd*CysZ R23A mutants all have severely impaired sulfate uptake capability, as demonstrated in cells and proteoliposomes (Fig. 5b, 4-S1a), consistent with a conserved role for these targeted arginine residues in all CysZ variants.

**Fig. 5.**
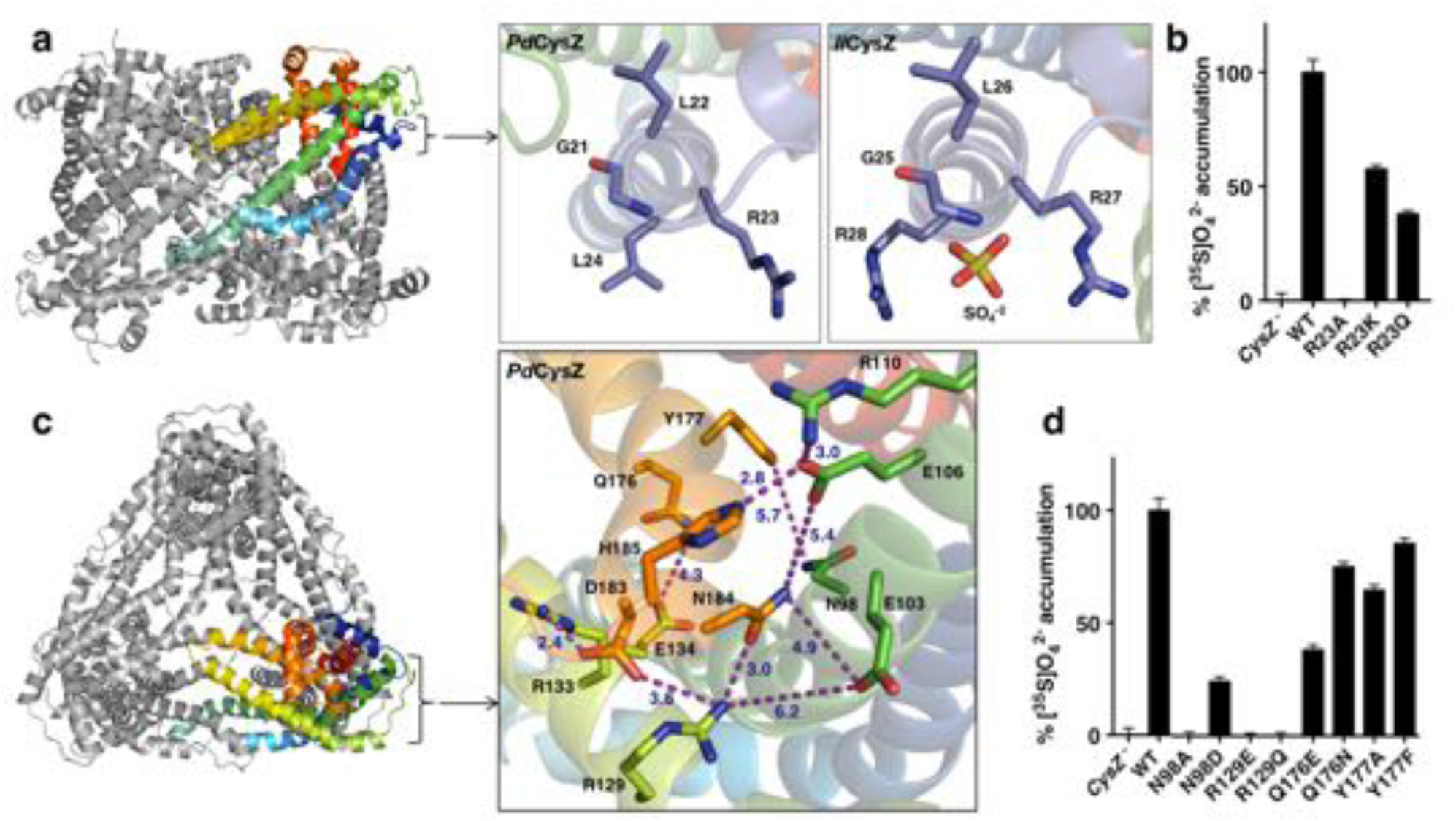
Functionally relevant features of the CysZ molecule. a. Side-view of a ribbon diagram of the *Pd*CysZ hexamer, with one chain rainbow-colored from N-(blue) to C-termini (red). Insets show the sulfate-binding site located at the start of H2a in *Pd*CysZ (left), with conserved residues G21, L22, R23 and L24 labeled, and the corresponding site in *Il*CysZ (right), with residues G25, L26, R27, and R28 labeled and showing the SO_4_ ^-2^ ion as bound in the crystal structure. b. Sulfate uptake by sulfate-binding site mutants of *Pd*CysZ. *E. coli K12* CysZ^-^ cells transformed with the listed *Pd*CysZ R23 mutants were used to measure [^35^S]O_4_ ^-2^ uptake (n=3). Sulfate uptake was abolished for R23A, and rescued to 50% and 40% of the wild type levels for the R23K and R23Q mutants respectively. c. A top-view of the *Pd*CysZ hexamer, colored as in a. The inset magnifies the central core of CysZ to show the associated network of hydrogen bonds (R129-N184, E106-H185), van der Waals interactions (E103-N184, E106-N184, Q176-E134) and salt bridges (R129-D183, R133-D183, R110-E106) between pairs of highly conserved residues. Interatomic contacts are shown as purple dotted lines with distances (in Å) marked in blue. d. Sulfate uptake by central-core mutants of *Pd*CysZ. *E. coli K12* CysZ^-^ cells transformed with the listed *Pd*CysZ mutants were used to measure [^35^S]O_4_ ^-2^ uptake (n=3). N98A and R129E and R129Q showed severely impaired sulfate uptake, whereas more conservative substitutions such as N98D, Q176E and Q176N had less of a negative effect on function. Y177A and Y177F do not show any impaired function.

### Conserved core in the hydrophilic head

Each apex of the triangular faces of the *Pd*CysZ hexamer comprises an extra-membranous hydrophilic head of a CysZ protomer (Fig. 1a and 5c). The central core of this hydrophilic head consists of the ends of H3 and H5 and the start of H4, with their residues forming an intricate network of hydrogen bonds and salt bridges that hold this helical bundle together (Fig. 5c). Two highly conserved motifs in this region, ExVE and QYxDYPxDNHK (Fig. 2c), likely play critical roles in this network of interactions. The two aspartates E103 and E106 (*Pd*CysZ) in the first motif interact with R129, N184 and R110, H185, respectively (Fig. 5c). The conserved R129 and R133 on H4b in turn also interact with D183 and N184 in the DNHK sequence stretch of H5b in the second motif. In the second motif, a conserved tyrosine (Y177) lies below the membrane interface towards the core of the molecule, along H5.

Mutational studies on a subset of these conserved hydrophilic-head residues highlight their functional relevance (Fig. 5d). R129 mutants (R129E or Q) exhibited a severe loss of SO_4_^2-^ binding and uptake, and mutations made to Q176 and Y177 also showed loss in function, albeit to a lesser extent. Charge reversal mutations made to E103 and E106 (to K or R) or the equivalent *Il*CysZ E107, E110 resulted in very poor expression and stability levels, suggesting that the disruption of the interaction network in this region may destroy the structural integrity of the protein.

### Putative pore and sulfate translocation pathway of CysZ

It was already evident from the initial *Il*CysZ structure that the hydrophilic head presented an incipient opening into the membrane, as seen in the center of the Fig. 5c inset, but it was quite unclear how this opening might relate to transmembrane sulfate translocation. The ‘transverse’ dimer structure of *Pf*CysZ clarified the possibility for ion permeation by showing that openings in its two protomers aligned across the putative membrane; however, the prospective transduction pathway would then be open on one side to the bilayer. The *Pd*CysZ structure showed that the open sides of transverse dimers line a central cavity in the *Pd*CysZ hexamers. Thus, plausible transduction pathways became more evident.

The view into a protomer surface along the direct pathway between transverse dimer apices (e.g. green to orange in Fig. 3e) displays the pattern of exceptionally high conservation associated with the entrance to a putative pore for sulfate translocation (Fig. 6a, b). This putative entrance or ‘pore’ lies in the midst of a tight network of interacting residues, which is seen detailed in the inset of Fig. 5c in the very same view as for Fig. 6a. These interactions close the incipient pore from ion conduction in this conformation. The electrostatic potential surface of PdCysZ (Fig. 6c) shows striking and puzzling electronegativity at the conserved pore entrance (Fig. 6d compared with Fig. 6b). Because of the conservations, similar electrostatics pertain to the two other homologs. Moreover, we observe the same helix dispositions, pore shape and charge interactions in all three structures, evident when the protomers are superimposed (Fig. 3c, top view).

**Fig. 6.**
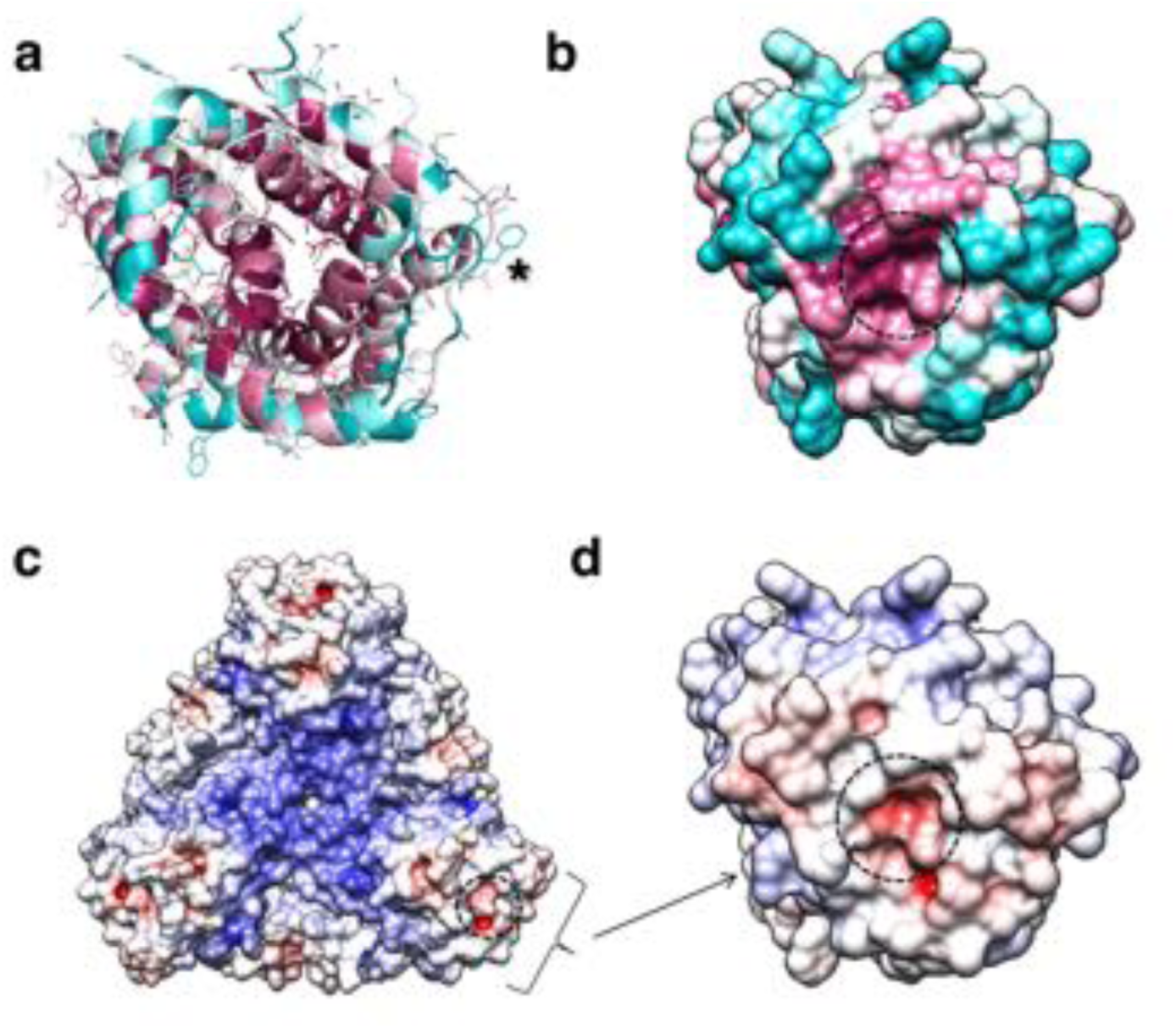
Conservation and entrance of putative pore of *Pd*CysZ. a. Ribbon diagram colored by conservation with residues in maroon being most conserved to cyan being least conserved (calculated by ConSurf) to highlight the entrance to the putative pore; an asterisk (*) marks the location of the sulfate-binding site (GLR motif) at top of H2a. b. Same view and coloring scheme as in a, but now shown in surface representation. c. Electrostatic representation of hexameric *Pd*CysZ as viewed from the top, with negative surface potential represented in red, and positive potential in blue as calculated by APBS, with location of the putative pore marked by a dashed circle. d. Close-up view in electrostatic representation of the putative pore within a *Pd*CysZ protomer, surface and orientation as in b.

Pathway prediction algorithms (namely, PoreWalker (Pellegrini-Calace, Maiwald, & Thornton, 2009)) performed on the CysZ protomer revealed a putative ion translocation pathway that begins at the entrance to the putative pore and ends in the large central cavity of the CysZ hexamer that lies within the membrane (Fig. 7a, b). In the structure of *Pd*CysZ, the entrance of the putative pore appears to be closed by the network of charge-interactions by conserved residues, R110, E106, N184, H185 and W235 (Fig. 6a, 7c-1). These residues interact to tightly restrict access to the putative translocation pathway and constrict the entry of ions, likely selected for size and charge (Fig. 7-S1a). After the narrow entrance, the putative pathway broadens, surrounded by a ring of conserved asparagine and glutamine residues, N33, N91, N98, Q176, and N220 (Fig. 7c-2). Mutations to N98 (N98A, N98D) led to impaired sulfate uptake (Fig. 5d). Following this polar environment, the putative pathway widens even further leading into a large, primarily hydrophobic, internal cavity encapsulated by the TM helices of the *Pd*CysZ hexamer, located in the plane of the lipid bilayer (Fig. 7b, 7-S1c, d). The hydrophobic cavity is enclosed on both the top and bottom by pairs of helices H4-H5, three on each side from the six protomers, pointing in towards the three-fold axis, with each side of the cavity, at its mid-section, measuring ∼50 Å, when viewed from above or outside the membrane (Fig. 7-S1c, d). Given the symmetric, dual topology nature of the CysZ assembly, the ions could then exit the cavity via the same pathway as they entered, but traveling through CysZ protomers located on the opposite side of the membrane.

**Fig. 7.**
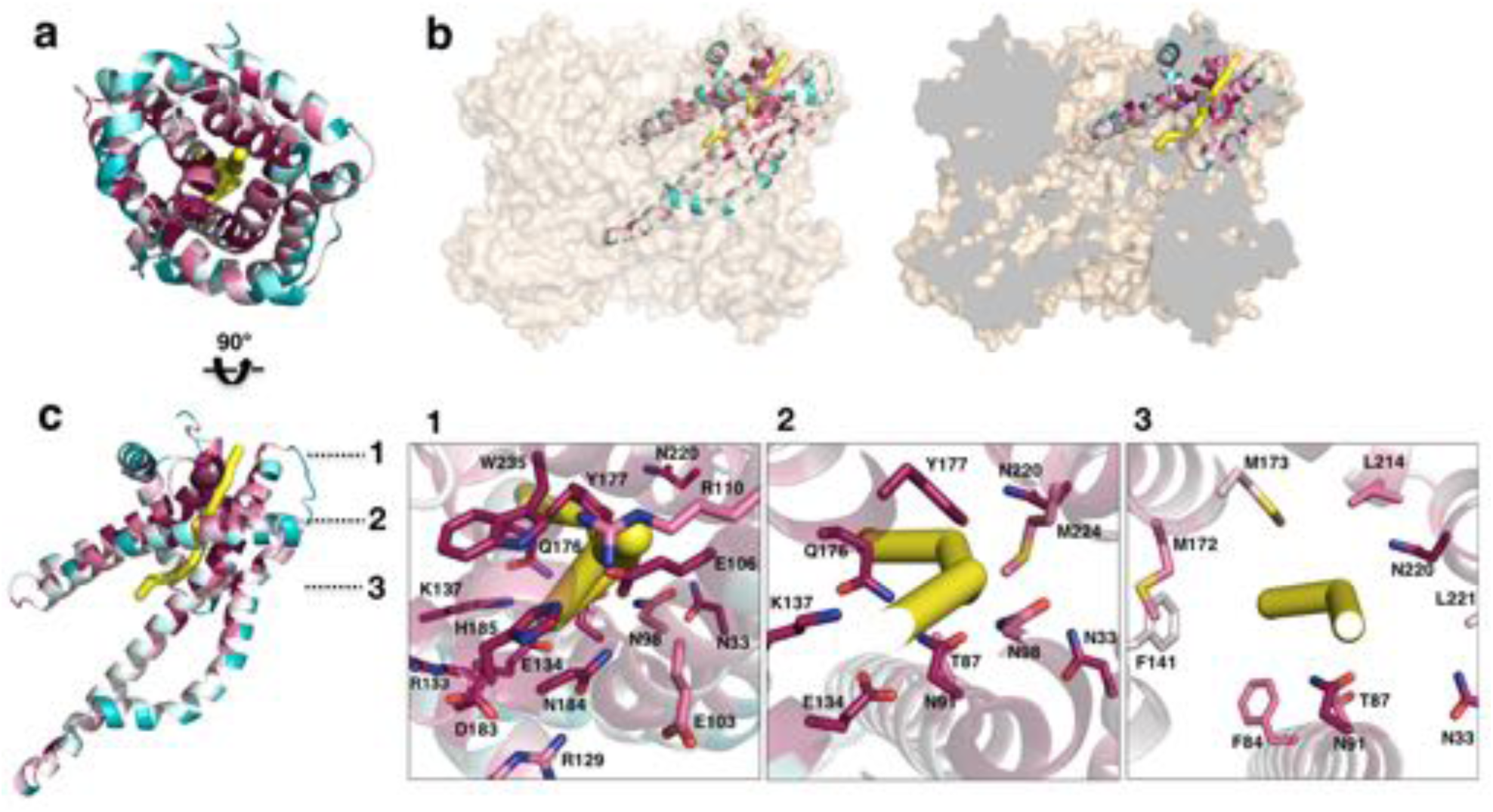
Putative ion conductance pathway of CysZ. a. A *Pd*CysZ protomer looking into the incipient pore entrance. The polypeptide ribbon is oriented as in 6a and colored by level of sequence conservation (calculated by ConSurf, with maroon being most conserved and cyan least). The putative pore and ion conduction pathway is shown as a yellow tube of 1 Å diameter, calculated by PoreWalker. b. Surface representation of the *Pd*CysZ hexamer, as side views, both semi-transparent (left) and with surface clipped (right, cut surfaces colored in grey) to allow for the internal visualization of the pathway leading to the central cavity. c. Side view of a *Pd*CysZ protomer left, viewed as in b and rotated 90° from a. Insets show magnified views of cross-sections along the pathway: At level 1, the entrance to the pore consists of a narrow constriction created by a network of highly conserved polar and charged residues (E106, R110, E134, N184, H185) tightly interacting with one another. At level 2, the pathway broadens and becomes less charged, but yet polar in nature as lined by conserved asparagine, tyrosine and threonine residues (N33, T87, N91, N98, Q176, Y177). From level 3, the pathway widens further and ultimately leads into the large central hydrophobic cavity.

## DISCUSSION

There are four known families of dedicated SO_4_^2-^ transport systems in prokaryotes^6^, of which CysZs are the least studied (Zhang et al., 2014). The sequence of CysZ, coding for an integral membrane protein with four predicted TM segments, shows no resemblance to any other known protein. This, and the quest to set the basis for a mechanistic understanding of function, prompted us to investigate the structure of CysZ.

We present here three structures of CysZ from different species, all determined by x-ray crystallography. These structures are all similar, showing a novel fold comprising two extended TM helices and two hemi-penetrating helical hairpins, giving rise to a tripod-like shape within the membrane, and a hydrophilic head (Fig. 2a, b, 3c). Two of the structures (*Il*CysZ and *Pf*CysZ) show dimers in the crystals, albeit with different interfaces (Fig. 3a, b), and the third structure (*Pd*CysZ) displays a hexameric assembly in multiple crystal forms (Fig. 1a).

All three structures have a dimeric component, each of which is present in the hexameric arrangement present in all the crystal forms of *Pd*CysZ (Fig. 3a, b), showing how they are all related. Furthermore, *Il*CysZ can be crosslinked in membranes to the alternative *Pf*CysZ dimer by engineering of disulfide cysteine mutants (Fig. 3-S2), which suggests that *Il*CysZ can adopt both of the dimer assemblies observed in the *Pd*CysZ hexamer. Thus, we hypothesize that CysZ is a hexamer in nature, as observed in *Pd*CysZ, and that in the case of *Il*CysZ and *Pf*CysZ, this assembly may have come apart into the subsequently crystallized dimeric components during the process of detergent extraction from the membrane and purification. This could be explained by the unusually labile, and conformationally-flexible, nature of the H4b-H5a helical hairpin of CysZ, with only one pair of stabilizing TM helices per protomer (H2-H3). We presume that the hexamer is maintained with the support and scaffolding of the lipid bilayer.

CysZ shows a clear dual topology arrangement, which we confirmed with cross-linking experiments on membranes (Fig. 3-S2). Dual-topology insertion is fairly uncommon in membrane proteins; however, reported cases include the well-documented EmrE, a multi-drug resistant export protein that inserts into the membrane as an anti-parallel dimer, and the more recently discovered and studied family of double-barreled ‘Fluc’ fluoride channels (Amadi, Koteiche, Mishra, & McHaourab, 2010; Korkhov & Tate, 2009; Rapp, Granseth, Seppala, & von Heijne, 2006; Stockbridge et al., 2015; Stockbridge, Robertson, Kolmakova-Partensky, & Miller, 2013).

Functional characterizations of the three CysZs for which we have obtained structural information show that all three mediate SO_4_^2-^ uptake and that this uptake is inhibited by SO_3_ ^2-^, for which the protein has higher affinity. Comparable results were obtained in cell-based as well as in proteoliposome-based uptake assays (Fig. 4b, d, 4-S1a). Affinities for SO_4_^2-^ and SO_3_ ^2-^ were measured by SPA (Fig. 4c). These results are consistent with previously reported data on *E. coli* CysZ (Zhang et al., 2014). Experiments performed on cells with uncoupling agents (Fig. 4-S1b) showed that CysZ-mediated SO_4_^2-^ uptake was independent of a classical electrochemical gradient, thus suggestive of the driving force being a concentration gradient of SO_4_^2-^ across the membrane. This hypothesis is supported by our single-channel bilayer experiments performed with *Pd*CysZ and *Il*CysZ, which show that CysZ has channel-like properties (Fig. 4-S2).

Building on our functional and structural discoveries, our results suggest a fascinating hypothesis for mechanisms of SO_4_^2-^ transfer and regulation. There is a conserved central core (Fig. 6a), which is likely to have a structural role. Close to this lies a sulfate-binding site, with functional implications (Fig. 5a), and the entrance to a putative pore (Fig. 5c). The entrance of this putative pore is delineated by a hydrophilic network of conserved residues that form a tight constriction in our observed conformation. Following this hypothetical route, sulfate ions that might enter through the three separate pores, one per CysZ dimer pair, would converge into a central hydrophobic cavity (Fig. 7b, 7-S1c, d). Once sulfate ions enter this central cavity, due to the unfavorable environment, these are likely to exit it rather rapidly through one of the three available exit pores on the cytoplasmic side of the hexamer. Hydrophobic cavities and pores are seen commonly in ion channels, with examples ranging from the well documented hydrophobic inner pores of the various potassium channels (Doyle et al., 1998) to the SLAC1 (Chen et al., 2010) and bestrophin anion channels (Yang et al., 2014), and the MscS and MscL mechanosensitive channels (Anishkin, Akitake, Kamaraju, Chiang, & Sukharev, 2010; Bass, Strop, Barclay, & Rees, 2002; Birkner, Poolman, & Kocer, 2012; Chen et al., 2010; Doyle et al., 1998), facilitating the rapid passage of ions due to the unfavorable environment, as well as potentially providing a means of ‘hydrophobic gating’ (Aryal, Sansom, & Tucker, 2015).

The surface electrostatics of CysZ (Fig. 1b and Fig. 6c, d) draw attention to the negative potential of the conserved core of each protomer, which is then surrounded by a more neutral annulus. While this feature seems contradictory to admission of SO_4_^2-^ ions at the extracellular side, it could be advantageous for expulsion into the cytoplasm on the opposite side. In any case, the structures that we have determined are evidently in a closed state, implying that a conformational change would have to occur to allow passage of SO_4_^2-^, likely modifying the surface electrostatics of the protein. It is tempting to speculate that the regulation of the opening of the pore could be modulated by the binding of sulfate ions to the identified sulfate-binding site, as it is near the entrance of the putative pore. L22 lies in proximity of the entrance of the putative pore, and its backbone amide (along with G21) coordinates the sulfate ion in the GLR motif of the sulfate-binding site. Thus, the binding of sulfate to the GLR motif could trigger a conformational change needed to displace L22, allowing for a wider opening for the sulfate ions to enter the pore. Sulfite could hypothetically exert its inhibitory effect on CysZ function by binding to this site.

There is an overall electropositive region in the center of the hexameric molecule, when viewed from the top (Fig. 1b, 6c, d). This central region is lined by conserved residues along helices H4b-H5a, namely, R129, R133 and K137. The hydrophobic tips of helices H4b-H5a of the three protomers on each side of the membrane then converge in the center of the hexamer. A pore through the three-fold axis of the hexamer could provide an alternative passageway for SO_4_^2-^ ions through this assembly. Although we cannot exclude it, this possibility seems less plausible because of the poor conservation at the tips of helices H4b-H5a and the strictly hydrophobic nature of this region.

We observe, in agreement with previous data, that SO_3_ ^2-^ inhibits CysZ-mediated SO_4_^2-^ flux (Fig. 4b, d, 4-S2a, f) (Zhang et al., 2014). The dual topology nature of CysZ could provide a means for internal as well as external regulation. However, experiments in solutions – such as crystallizations – where all molecules are exposed to the same chemical environment, makes capturing such a state in an open or SO_3_ ^2-^ blocked conformation challenging. Despite this limitation, our CysZ structures and associated functional experiments have allowed us to make substantial progress in the understanding SO_4_^2-^ uptake by these membrane permeases. This work sets the framework for future experiments aimed at unraveling the molecular details of how SO_4_^2-^ is translocated across the membrane by CysZ and how this process is regulated.

### PDB Accession codes

*Idiomarina loiheinsis* CysZ: **3TX3**, *Pseudomonas fragi* CysZ: **TBD,** *Pseudomonas denitrificans* CysZ: **TBD**.

## Materials and Methods

### Ortholog selection and cloning

A total of 63 *cysZ* candidate genes were selected by a bioinformatics approach implemented by the New York Consortium of Membrane Protein Structure (NYCOMPS), as previously described (Punta et al., 2009). The majority of the genes (including *Il*CysZ, uniprot ID: Q5QUJ8) were PCR-amplified from fully sequenced prokaryotic genomic DNA (obtained from ATCC^®^) (Love et al., 2010). CysZ genes from certain species, such as *Pf*CysZ (uniprot ID: A0A0X8F058) and *Pd*CysZ (uniprot ID: M4XKU7) were chemically synthesized by GenScript^®^, with codon-optimization for protein expression. All genes were cloned by ligation-independent cloning (LIC) (Aslanidis & de Jong, 1990) into an IPTG (isopropyl β-D-1-thiogalactopyranoside) inducible, kanamycin-resistant pET derived plasmid (Novagen), with an N-terminal decahistidine tag (His10) and a TEV (tobacco etch virus) protease site to allow for tag cleavage upon purification.

### Protein expression and purification

Expression plasmids bearing the *cysZ* genes were transformed into BL21(DE3)pLysS cells using standard protocols, and grown at 37 °C in 2XYT media supplemented with 50 μg/ml kanamycin and 50 μg/ml chloramphenicol in an orbital shaker at 250 rpm. Protein expression was induced for ∼16 hours at 22 °C with 0.2 mM IPTG once an absorbance (A_600 nm_) of 0.8-1.0 was reached. Selenomethione (Se-Met)-incorporated proteins were expressed in BL21(DE3)pLysS cells grown using an M9 minimal media kit (Shanghai Medicilon Inc.) supplemented with the necessary minerals, vitamins and non-inhibitory amino acids. Se-Met was added prior to IPTG induction at an A_600 nm_ of 1.2. The Se-Met-incorporated protein was purified using the same procedures as the native protein. Once harvested, the cells were resuspended at 0.2 g/ml in lysis buffer containing 20 mM Na-Hepes pH 7.5, 200 mM NaCl, 20 mM MgSO4, DNase I and RNase A, 0.5 mM PMSF (phenylmethylsulfonyl fluoride), EDTA-free Complete protease inhibitor cocktail (Roche) and 1 mM TCEP-HCl (Tris (2-carboxyethyl) phosphine hydrochloride) as a reducing agent. Initial small-scale expression and detergent screening was performed on 80 mg of pelleted cells (wet weight), and 7-10 g of cells for large-scale protein purification. Cells were lysed using an Avestin^®^ EmusiFlex-C3 homogenizer, followed by protein solubilization with 1% (w/v) decyl maltopyranoside (DM) (Anatrace^®^ Affymetrix) for 1 hour at 4 °C, after-which insoluble material was removed by ultra-centrifugation at 100,000 x g. The solubilized protein was applied to Ni-NTA Sepharose (Qiagen) in batch, washed with lysis buffer containing 0.2% DM and 40 mM imidazole and eluted in buffer containing 250 mM imidazole. Upon elution, CysZ was dialyzed overnight with His-tagged TEV protease at 4°C, against a buffer containing 20 mM Na-Hepes pH 7.0, 200 mM NaCl, 0.2% DM, 1 mM TCEP-HCl and 20 mM Na2SO4, allowing for the cleavage of the His10 tag and removal of the imidazole. Tagless CysZ was then re-passaged over Ni-NTA sepharose to re-bind of any uncleaved CysZ, TEV protease and the cleaved His10 tag. The protein was then subjected to size-exclusion chromatography (Superdex 200 10/30 HR, GE Healthcare) in 20 mM Na-Hepes pH 7.0, 200 mM NaCl, 1 mM TCEP-HCl, 20 mM Na2SO4 and appropriate detergent for crystallization, for *Il*CysZ: 0.06% Lauryl dimethylamine oxide (LDAO) and for *Pf*CysZ and *Pd*CysZ: 1% β-octyl glucopyranoside (β-OG). The choice of detergent was made based on protein yield, stability and mono-disperse gelfiltration peaks obtained in the initial small-scale detergent screening. A yield of ∼1.5 mg of purified CysZ was typically obtained from a cell pellet of 7-8 grams (1 liter of culture).

### Protein Crystallization

*Il*CysZ: Crystals of *Il*CysZ in LDAO were obtained by vapor diffusion at a protein concentration of 6-8 mg/ml at 4°C, in a 1:1 v/v ratio against a precipitant of 28-32% PEG400, 0.1M Tris-HCl pH 8.0, with salt additive of 0.1 M NaCl or 0.1 M MgCl_2_. The crystals appeared overnight, continued to grow in size over the course of 2-4 days after set-up. After optimization, the crystals grew to a maximum size of ∼200 μm x 100 μm x 50 μm with a rhomboid or cuboid shape. The crystals were harvested directly without the addition of a cryo-protectant and flash-frozen into liquid nitrogen, for data collection on the X4A/X4C beamlines at National Synchrotron Light Source (NSLS), Brookhaven National Labs (Upton, NY). SeMet derivatized and selenate cocrystals were obtained from the same conditions as the native crystals.

*Pf*CysZ: Crystals of *Pf*CysZ in β-OG at 5 mg/ml were initially obtained at 4 °C by vapor diffusion, in a 1:1 protein to precipitant ratio, after 1-2 days against 28% PEG400, 0.1 M MES pH 6.0. They were cuboid in shape and grew in clusters of multiple crystals originating from a common locus. The crystals were optimized to a maximal size of ∼150-200 μm x 50 μm x 50 μm, with the best diffracting crystals grown under silicone oil (visc. 500) in microbatch Terazaki plates. Crystals were directly flash-frozen into liquid nitrogen without the use of a cryo-protectant and were exposed to X-rays at the NE-CAT (241DC and E) beamlines at APS, Argonne National Lab (Argonne, IL) for data collection. SeMet crystals were obtained from the same conditions as the native protein.

*Pd*CysZ: Crystals of *Pd*CysZ in β-OG at 5-8 mg/ml were initially obtained at 4°C by vapor diffusion, in a 1:1 protein to precipitant ratio, after 2-3 days against 22-30% PEG550MME, 0.1 M Na-Hepes pH 7.0. The rod-like crystals were hexagonal on one face, and grew in clusters originating from a common locus as well as on the edge of the drop. The multiple crystal forms observed were all obtained in the same crystallization conditions. The crystals were optimized to a maximal size of ∼200-250 μm x 25 μm x 50 μm. With the addition of 20% glycerol (w/v) as a cryo-protectant, the crystals were flash-frozen into liquid nitrogen and were exposed to X-rays at the NE-CAT (24-IDC and 24-IDE) beamlines at APS, Argonne National Lab (Argonne, IL) for data collection.

### Data collection and structure determination

*Il*CysZ: The structure of CysZ was determined by the single-wavelength anomalous diffraction (SAD) method from anomalous diffraction of a selenate (SeO_4_^2-^) derivative crystal. The anomalous signals were measured at the Se K-edge peak wavelength, which was determined experimentally from fluorescence scanning of the crystal prior to data collection. All diffraction data were recorded at 100K using an ADSC Q4R CCD detector at the NSLS X4 beamline. Diffraction data were indexed, integrated, scaled, and merged by HKL2000 (Otwinowski & Minor, 1997). Selenate substructure determination was performed with the SHELXD program through HKL2MAP (Pape & Schneider, 2004). A resolution cut-off at 2.6 Å was used for finding Se sites by SHELXD. A strong peak found by SHELXD was used to calculate initial SAD phases, which were improved by density modification by SHELXE (Sheldrick, 2010). With a solvent content of 65% corresponding to 2 molecules in the asymmetric unit, 50 cycles of density modification resulted in an electron density map of sufficient quality for model building. The initial polypeptide chain was built by Arp/Warp (Langer, Cohen, Lamzin, & Perrakis, 2008), at 2.1 Å by using experimental phases. Further cycles of model building were performed manually using COOT (Emsley, Lohkamp, Scott, & Cowtan, 2010) and all rounds of refinements were performed with PHENIX (Adams et al., 2010). The native structure of CysZ with bound sulfate was determined both by multi-crystal native SAD (Liu et al., 2012) (final resolution of 2.3 Å) and by molecular replacement with the selenate bound model (final resolution of 2.1 Å). In addition, phase information obtained from Se-Met derivatized protein with 9 Se sites per CysZ molecule, verified our model obtained from the selenate data.

*Pf*CysZ: Multi-crystal SeMet-SAD data sets were collected at APS beamline 24-IDC with a Pilatus 6M pixel array detector under a cryogenic temperature of 100 K. To enhance anomalous signals from Se atoms for phasing (Liu, Zhang, & Hendrickson, 2011), the X-ray wavelength was tuned to the Se-K edge (λ=0.9789 Å). The orientation of crystals was random without special consideration of crystal alignment, and beam size was adjusted to match the crystal size. A total of 22 data sets were collected, each from a single crystal. An oscillation angle of 1° was used for data collection with a total of 360 frames for each data set. The beam size was adjusted to match the crystal size. The 22 single-crystal data sets were processed individually by using XDS (Kabsch, 2010) and CCP4 packages (Winn et al., 2011). For phasing purposes, the low-resolution anomalous signals were enhanced by increased multiplicity. By rejection of 7 outlier crystals (Liu et al., 2012), anomalous diffraction data from 15 statistically-compatible crystals were scaled and merged for phasing. For outlier rejection, a unit-cell variation of 1.0σ was used. CCP4 program POINTLESS and SCALA (Evans, 2006) were used for data combining; and Bijvoet pairs were kept separately throughout the data flow. For refinement purposes, keeping the high angle data was important, and done by limiting radiation damage as well as by increasing multiplicity. Although most *Pf*CysZ crystals diffracted to only about 3.5 Å spacings or poorer, we intentionally set the detector distance to include higher spacings. Higher resolution data were retained through a data merging procedure that is described as follows: 1) The 22 individually processed data sets were analyzed by diffraction dissimilarity analysis by using only high angle data between 3.5-3.0 Å, resulting in three subsets. 2) The data statistics of members in each subset were checked manually and the subset that contained the highest angle data set, e.g. data set 6, was selected for further analyses and data combination. 3) Each data set within the selected subset was compared with data set 6 by high-angle intensity correlation. Six of the highest resolution data sets were statistically comparable and therefore were selected for merging. For phasing, substructure solutions were found by SHELXD (Sheldrick, 2010) and were further refined and completed by PHASER (McCoy et al., 2007) and then used to compute initial SAD phases at the data limit by SAD phasing with PHENIX (Adams et al., 2010). Phases were density modified with solvent flattening and histogram matching as implemented in CCP4 program DM (Cowtan & Zhang, 1999) to improve phases and also to break phase ambiguity. The estimated solvent contents of 71% were used for density modification. The model was initially built into the experimental electron density map by COOT (Emsley et al., 2010), followed by iterative refinement by PHENIX and model building in COOT. The refined model does not contain solvent molecules at this resolution.

*Pd*CysZ:

Native crystal data were collected at the APS beamline 24-IDC with a Pilatus 6M pixel array detector at a cryogenic temperature of 100 K at an X-ray wavelength of λ=1.023 Å. The sample-to-detector distance was set to 500 mm. An oscillation angle of 0.5° was used for data collection. The beam size was adjusted to match the crystal size. Molecular replacement was attempted using a variety of search models (*Pf*CysZ and *Il*CysZ monomer/dimer models, with various degrees of truncation). Success was achieved by searching for six copies of a search model consisting of the *Pf*CysZ monomer, with residues 36-53 deleted and the sequence adjusted using CHAINSAW (Stein, 2008) (pruning non-conserved residues to the gamma carbon). Density modification of the initial map was performed in PARROT, incorporating solvent flattening, histogram matching and NCS-averaging. An initial round of model building was performed in COOT (Emsley et al., 2010) into this map, followed by further phase improvement and bias-removal using *phenix.prime_and_switch* (Adams et al., 2010). The improved map was used for a second round of model building, followed by iterative cycles of reciprocal space refinement using *phenix.refine*, and real-space refinement and correction in COOT.

All graphical representations and figures of our structural models were made in either PyMOL (Schrodinger, 2010) or Chimera (Pettersen et al., 2004).

### Radioligand Binding by Scintillation Proximity Assay (SPA)

CysZ was purified by standard protocols as described above, with the exception of leaving the histidine-tag intact without cleavage by the TEV protease. The imidazole was removed by dialysis and the purified protein was not run over the size exclusion column, and instead was directly concentrated to 2 mg/ml for the SPA experiment. [^35^S]O_4_^2-^ obtained in the form of sulfuric acid (American Radiolabeled Chemicals (ARC)) was used as the radioligand. 100 ng – 250 ng of CysZ was used per assay point, diluted in 100 μl of assay buffer containing 20 mM Hepes pH 7.0, 200 mM NaCl, 0.2% DeM, 20% glycerol, 0.5 mM TCEP, with 12.5 μl of Copper YSi beads (Perkin Elmer). A trace amount of [^35^S]O_4_^2-^ (∼10-30 nM) mixed with cold ligand (Na_2_SO_4_) was added to each well of a 96 well clear bottom plate along with the mix of protein, buffer and beads. The plate was agitated for a minimum of 30 minutes to as long as overnight, and measured in a scintillation counter the following morning (MicroBeta, Perkin Elmer). Competitive binding experiments with a gradient of a competing cold ligand ranging from (10 nM – 100 mM) were performed in triplicate, with the control measured at each concentration by adding 1 M Imidazole to the reaction buffer, to prevent binding of the protein to the scintillation beads. All data were analyzed and graphically represented with GraphPad™ Prism6 software.

### [^35^S]O_4_^2-^ Uptake Experiments

*In whole cells:* The *cysZ* gene knockout strain in *E. coli* K-12 was obtained from the Coli Genetic Stock Center at Yale University (http://cgsc.biology.yale.edu/), originally found in the Keio knockout collection (Baba et al., 2006). Genotype: F-, β, (araD-araB)567, β, lacZ4787(::rrnB-3), *λ* -, β, cysZ742::kan,rph-1, β, (rhaD-rhaB)568, hsdR514. The knockout strain was made competent by standard protocols (Hanahan, 1983) to allow for the transformation and expression of *cysZ*-containing plasmids, for rescue experiments. Wild-type parental *E. coli* K-12 was used as a control strain. The strains were grown in LB (Luria Broth) without any antibiotic (for the WT cells) and with 50 μg/ml of kanamycin for the knockout cells at 37 °C overnight to saturation. The culture was spun down at 3000 x g the next morning and the cells were resuspended in Davis-Mingioli (DM) minimal media without sulfate (MgSO_4_ was replaced with MgCl_2_ and (NH_4_)_2_SO_4_ was replaced by NH_4_Cl), supplemented with 0.63 mM L-cysteine to allow the cells to grow in the absence of sulfate (Davis & Mingioli, 1950). The cells were incubated in DM media for a minimum of 9 hours, maximum overnight, at 37°C after which they were spun down and washed 3 times in 5 mM Na-Hepes pH 7.0. This ensured that the cells were starved of SO_4_, to deplete the sulfate stores in the cell, enhancing sulfate uptake measured (F. Parra, personal communication). The cells were resuspended in the same buffer at 0.7 g/ml at room temperature, and 10 μl of the cell suspension was used for each point in the uptake experiment. A final concentration of 320 μM of SO_4_ (Na_2_SO_4_ with [^35^S]O_4_^2-^ as a tracer) was used outside. The uptake experiments were performed in triplicate, measured over a time course of typically 0 - 300 seconds. The reaction was stopped by dilution, by adding 1.5 ml of ice-cold buffer (5 mM Na-Hepes pH 7.0) to the cells in the tube. The reaction mix was immediately poured onto a vacuum filtration device, with an individual glass-fiber filter (0.75 μm, GF/F) per assay point, after which the filters are washed once more with 1.5 ml of buffer. The dry filters, with the cells attached to their surface, were then moved to scintillation vials, containing 4 ml of EconoSafe^®^ scintillation cocktail, to be counted the next morning. For the *CysZ*^*-*^ cell rescue experiments, a similar protocol was used, transformed with an ampicillin resistant expression vector containing the CysZ gene, mutant or empty vector (as the control). Upon transformation, the cells were grown overnight in a 4 ml starter culture of LB with 50 μg/ml kanamycin and 100 μg/ml ampicillin. The next morning the entire 4 ml was used to inoculate 75 ml of LB (Kan, Amp), and grown at 37 °C for 2.5-3 hours until an OD_600_ of 0.6-0.8 was reached. Protein expression was then induced at 37 °C for 4 hrs with 0.2 mM IPTG. After 4hrs, the cells were spun down and resuspended in the DM minimal media without sulfate, 0.63 mM cysteine, Amp, Kan and 0.2 mM IPTG, and incubated overnight at 22 °C. The next morning the cells were spun down, washed 3 times with 5 mM Na-Hepes 7.0 and resuspended at final concentration of 0.7 g/ml for uptake.

*In proteoliposomes:* CysZ was purified by the procedure described above, and upon elution from the size-exclusion column concentrated to 1 mg/ml for reconstitution into liposomes at a protein to lipid ratio of 1:100. The liposomes were comprised of a 3:1 ratio of *E. coli* polar lipids and phosphatidylcholine (PC) (Avanti Polar Lipids), prepared by previously described methods (Rigaud, Pitard, & Levy, 1995). Typically, 10 μl of CysZ-proteoliposomes and control ‘empty’ liposomes at a concentration of 5-10 mg/ml, were used per assay point and diluted into a reaction buffer of 100 μl, containing [^35^S]O_4_^2-^ (at trace amounts of ∼10-15 nM) supplemented with Na_2_SO_4_ at the desired concentration. The experiment was performed in triplicate and sulfate accumulation was measured either over a time-course or at a fixed time with different amounts of SO_4_^2-^ or SO_3_ ^2-^ present. The reaction was stopped by dilution, by adding 1.5 ml of ice-cold buffer (5 mM Na-Hepes pH 7.0) to the cells in the tube. The reaction mix was immediately poured onto a vacuum filtration device, with an individual nitrocellulose filter (0.22 μm) per assay point, after which the filters are washed once more with 1.5 ml of buffer. The dry filters, with the cells attached to their surface, were then moved to scintillation vials, containing 4 ml of EconoSafe^®^ scintillation cocktail, to be counted the next morning.

#### Measurement of CysZ single-channel activity in the planar lipid bilayer

A previously described method (Mueller, Rudin, Tien, & Wescott, 1962) to insert ion channels incorporated in liposomes into “painted” planar lipid bilayers by vesicle fusion was used to incorporate CysZ into a lipid bilayer created on a small aperture between two aqueous compartments, called the cis and trans compartments (Morera, Vargas, Gonzalez, Rosenmann, & Latorre, 2007). Since this system is very sensitive to contaminants, CysZ was expressed in and purified from a porin-deficient strain of *E. coli* cells to prevent any carry through of contaminating porins that could create large conductances and artifacts in the single-channel recordings. Phosphatidylethanolamine (PE) and phosphatidylserine (PS) (Avanti Polar Lipids), in a ratio of 1:1 were dissolved in chloroform to mix, and dried completely under an argon stream. The mixed and dried lipids were then dissolved in n-decane to a final concentration of 50 μg/ml, and kept at 4 °C. The lipids are always prepared fresh, on the same day of the experiment. The purified CysZ protein were incorporated into PE:PS (1:1) liposomes by brief sonication at 80kHz for 1 minute at 4 °C. The experimental apparatus consisted of two 1-ml buffer chambers separated by a Teflon film that contains a single 20- to 50-μm hole. A lipid bilayer was formed by “painting” the hole with the 1:1 mixture of PE:PS, this results in a seal between the two cups formed by the lipids (Leal-Pinto, London, Knorr, & Abramson, 1995). For these studies, the cis side was defined as the chamber connected to the voltage-holding electrode and all voltages are referenced to the trans (ground) chamber. Stability of the bilayer was determined by clamping voltage at various levels. If a resistance > 100 Ω and noise < 0.2 pA were maintained in the patch, the proteoliposomes containing CysZ were added to the trans side of the chamber and stirred for 1 min. The fusion event or insertion of a channel into the bilayer was assessed by the presence of clear transitions from 0 current to an open state.

### Site-specific cysteine labeling experiments

All functional and cysteine mutants of CysZ were generated by site-directed mutagenesis, using the QuikChange^®^ site-directed mutagenesis kit (Strategene). The sequence-verified mutants were then tested for expression in comparison to the WT CysZ. To address the membrane topology of *Il*CysZ, single cysteine mutants were designed to perform site-directed fluorescence labeling based on the accessibility of the cysteine to the membrane impermeable thiol-directed fluorescent probe (Ye, Jia, Jung, & Maloney, 2001). A set of surface-exposed residues at different positions on the CysZ molecule were selected to be mutated to cysteines, based on the *Il*CysZ structure. The cysteine mutants were expressed by standard protocols that were used for the WT protein. The membrane fraction of each mutant was pelleted after cell lysis by ultracentrifugation at 100,000 x g and resuspended at 20 mg/ml (Bradford assay) in fresh buffer containing 20 mM Na-Hepes pH 7.0, 200 mM NaCl, protease inhibitors: 0.5 mM PMSF and Complete protease inhibitor cocktail EDTA-free and 1 mM TCEP-HCl. 1 ml of membranes were then incubated with 30 μM membrane-impermeant fluorescein-5-maleimide dye (Invitrogen, 2 mM stock freshly prepared in water, protected from light) for 30 minutes in the dark at room temperature. The labeling reaction was stopped by the addition of 6 mM β-mercaptoethanol, and the membranes were spun down and resuspended in fresh buffer to remove any remaining unreacted fluorescent dye. The fluorescently labeled protein was then purified from the membranes by solubilization with 1% DM, using a standard Ni-NTA purification protocol, qualitatively analyzed on an SDS-PAGE, and quantitatively measured by a Tecan^®^ fluorescence plate reader at an excitation of 495 nm, emission of 535 nm.

### Crosslinking of Cysteine Mutant Pairs in CysZ

Cysteine mutants of *Pf*CysZ and *Il*CysZ were designed based on pairs of residues that were in close proximity (within 3-7 Å) of each other in our structure that could have the ability to covalently join the 2 protomers of the dimer. The single and double cysteine mutants were made by site-directed mutagenesis using the QuikChange^®^ Site-Directed Mutagenesis Kit (Strategene). Mutants and WT were expressed using the standard protocols CysZ expression. Isolated membrane fraction was resuspended by homogenization in fresh lysis buffer at a membrane protein concentration of ∼25 mg/ml (measured by Bradford Assay). Bis-methanethiosulfonate (Bis-MTS) crosslinkers (Santa Cruz Biotechnolgies) of different spacer lengths were used at 0.5mM (dissolved in DMSO) added to 1ml of resuspended membranes at RT for 1 hour. Bis-MTS crosslinkers are membrane permeable, highly reactive and specific to sulfhydryl groups, and the covalent linkage is resistant to reducing agents like β-mercaptoethanol (Akabas, Stauffer, Xu, & Karlin, 1992). The crosslinking lengths used were: 1,1-Methanediyl Bismethanethiosulfonate (3.6 Å), 1,2-Ethanediyl Bismethanethiosulfonate (5.2 Å), 1,4-Butanediyl Bismethanethiosulfonate (7.8 Å) and 1,6-Hexanediyl Bismethanethiosulfonate (10.4 Å). The reaction was quenched by the addition of 10 mM free cysteine, and the protein was then extracted and purified from the membranes with 1% DM followed by Ni-NTA resin. The imidazole is then diluted out in the Ni-elute, and the His-tag is cut with TEV protease in small scale. The cleaved protein is passaged over Ni-NTA resin to remove contaminants and uncleaved protein, and flowthrough is run on a reducing SDS-PAGE and stained with Coomassie blue to analyze dimer formation. The mutants designed were L161C (single mutant), L161C-A164C and N160C-L168C (double mutants) for *Pf*CysZ; and L156C-Q163C and V157C-Q163C (double mutants) for *Il*CysZ.

## Acknowledgments

Crystallographic data for this study were collected on the NSLS beamline X4A at Brookhaven National Laboratory and on the NE-CAT beamlines 24ID-C and E (supported by NIH-NIGMS grant P41 GM103403) at the Advanced Photon Source. The Pilatus 6M detector on 24-ID-C beam line is funded by a NIH-ORIP HEI grant (S10 RR029205). This work was supported by NIH-NIGMS grants R01 GM098617 (F.M.) and R01 GM107462 (W.A.H.), and by grants through the New York Structural Biology Center for the New York Consortium on Membrane Protein Structure (NYCOMPS; U54 GM095315) and for the Center on Membrane Protein Production and Analysis (COMPPÅ; P41 GM116799). O.B.C. was supported by a Charles H. Revson Senior Fellowship. We thank John Schwanof for assistance during data collection at NSLS, Joe Mindell, Francisco Parra, Albano Meli, Giuliano Sciara and Carlos A Villalba Galea for helpful advice and contributions at the early stages of the project, and Leora Hamberger for her assistance managing the Mancia Lab.

### Declaration of Competing Interests

The authors declare that they have no financial or non-financial competing interests.

---

**Fig. 1. S1.**
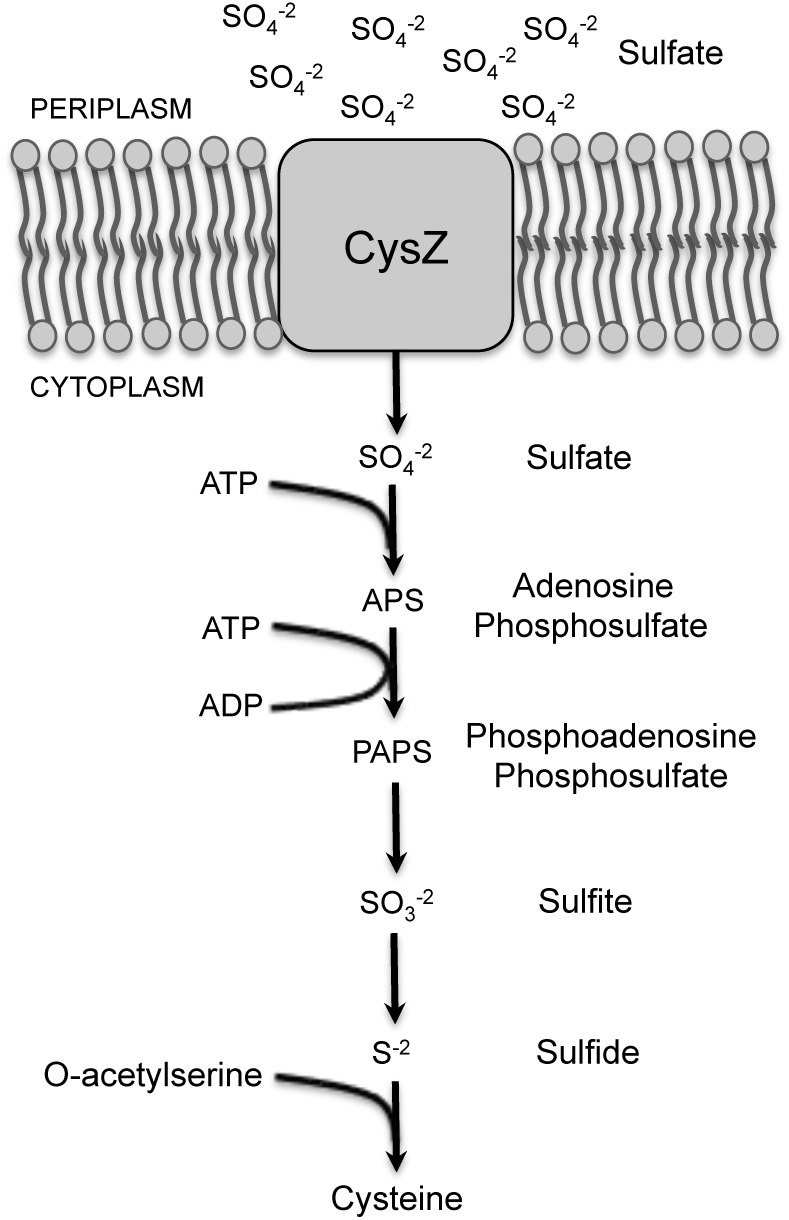
Schematic of assimilatory sulfate reduction in bacteria for cysteine biosynthesis. Assimilatory reduction is energetically dispendious as sulfate is extremely stable. In this pathway, sulfate ions enter the cell via a transporter or channel, like CysZ; this is followed by its activation by ATP-sulfurylase, which utilizes ATP to form adensosine phosphosulfate (APS) and inorganic pyrophosphate (PPi) as a by-product. APS is then further activated by the addition of a second phosphate forming phosphoadenosine phosphosulfate (PAPS) and this high-energy intermediate product is poised for sulfur removal to form sulfite (SO3^2-^) by the thioredoxin enzyme, which is then further reduced to sulfide ions (S^-^). Sulfide, the final reduced sulfur product, can then be used in a variety of ways by the cell, such as the incorporation into O-acetylserine to synthesize cysteine.

**Fig. 1. S2.**
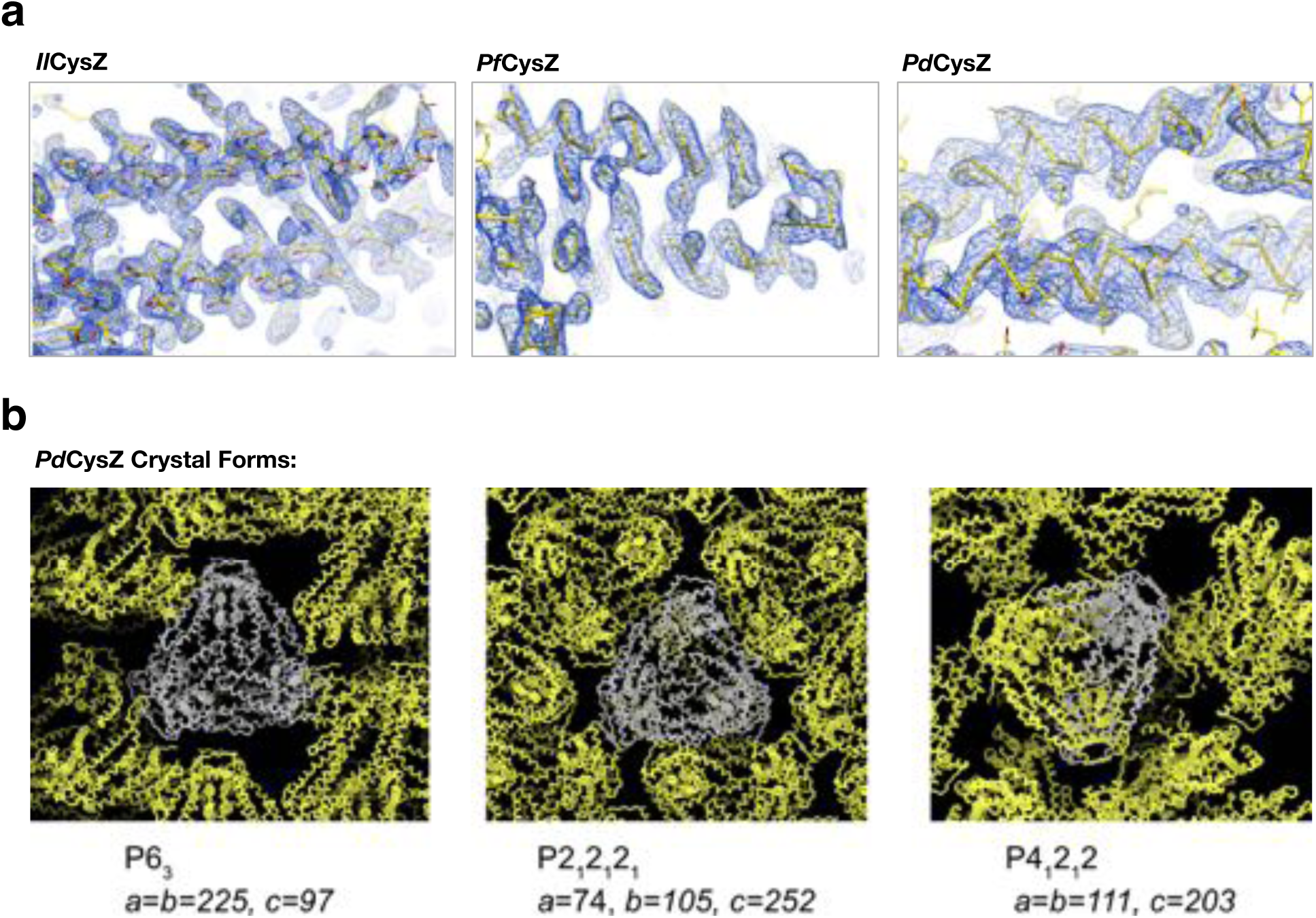
Representative electron density of the crystal structures of *Il*CysZ, *Pf*CysZ and *Pd*CysZ, and the different crystal forms observed for *Pd*CysZ. a. Representative electron density in blue mesh for: *Il*CysZ, contoured at 1 x r.m.s.d., *Pf*CysZ, contoured at 2 x r.m.s.d, with a negative B-factor of −100 Å^2^ applied to map; *Pd*CysZ, contoured at 1 x r.m.s.d, with a negative B-factor of −100 Å^2^ applied to map. Final atomic models (Cα trace with side chains depicted) are shown as yellow stick representation. b. Preliminary molecular replacement analyses of the 3 different crystal forms obtained for *Pd*CysZ, showing the hexameric assembly of the molecule, seen in each case. The Cα trace of an asymmetric unit in each case is depicted in grey, with their symmetry mates are shown in yellow. Space group and unit cell dimensions are listed below.

**Fig. 3. S1.**
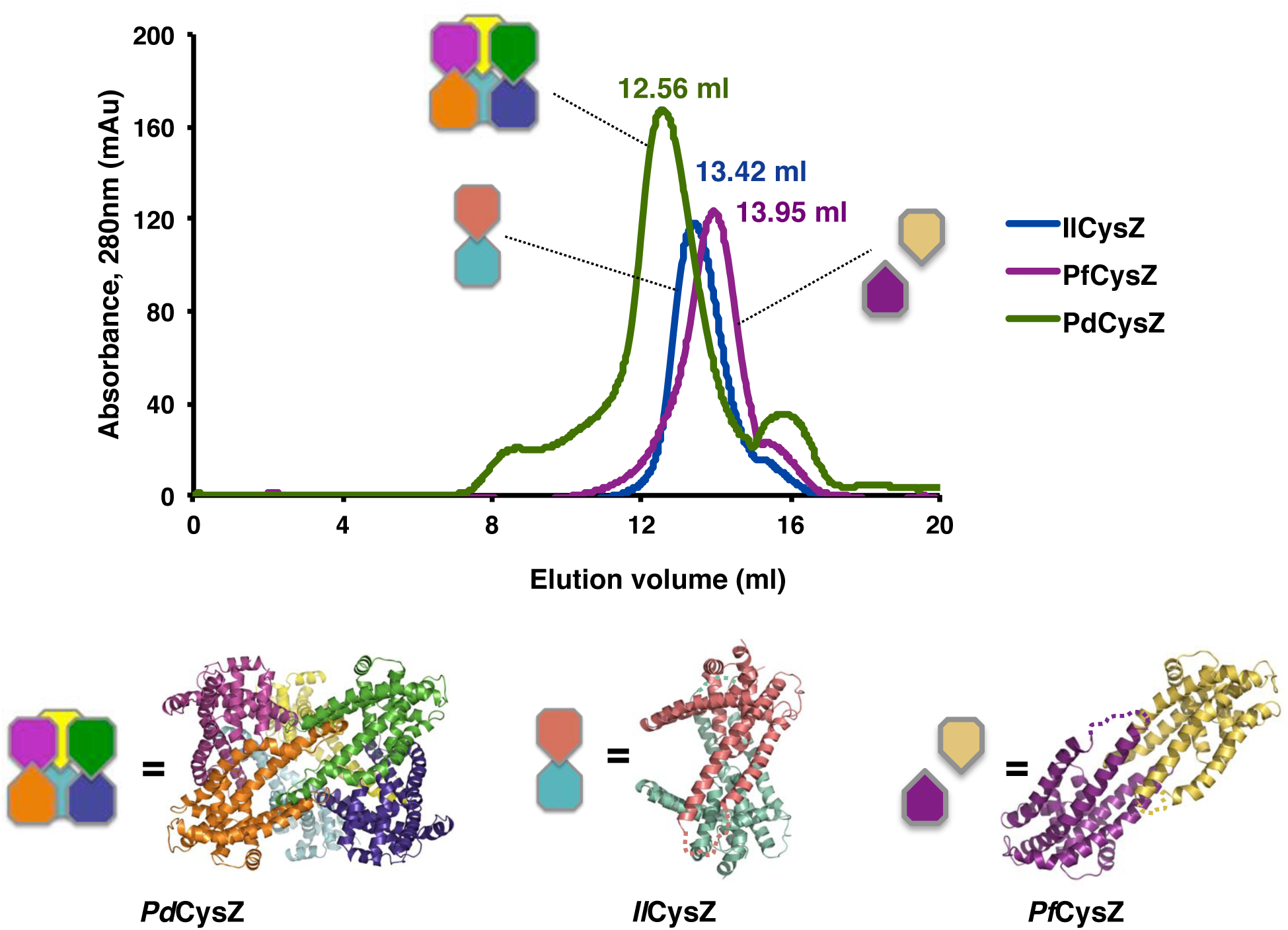
Size-exclusion chromatography of CysZ shows a mono-disperse elution profile for each of the three species purified - *Pd*CysZ, *Pf*CyZ and *Il*CysZ. All three proteins were solubilized from isolated membrane fractions purified in the presence of 0.2% decyl maltopyranoside (DeM) and exchanged into buffer containing 1% β-octyl glucopyranoside (β-OG) on the size-exclusion column (Superdex 200 10/30 HR). A schematic of the oligomeric states observed in the three crystal structures of CysZ are shown below the elution profiles. Crystallization trials were set-up directly after the size-exclusion chromatography step. A shift of ∼1 ml was observed for the elution of *Pd*CysZ (peak at 12.56 ml), with respect to *Pf*CysZ and *Il*CysZ, indicating that the size (volume) and shape of *Pd*CysZ is significantly larger than the other two species. *Il*CysZ elutes at 13.42 ml, 0.5 ml ahead of *Pf*CysZ, which could be explained by the shape of *Il*CysZ, occupying more volume as compared to *Pf*CysZ, with its helices (H4-H5) pointing outward, away from the body of the dimer.

**Fig. 3. S2.**
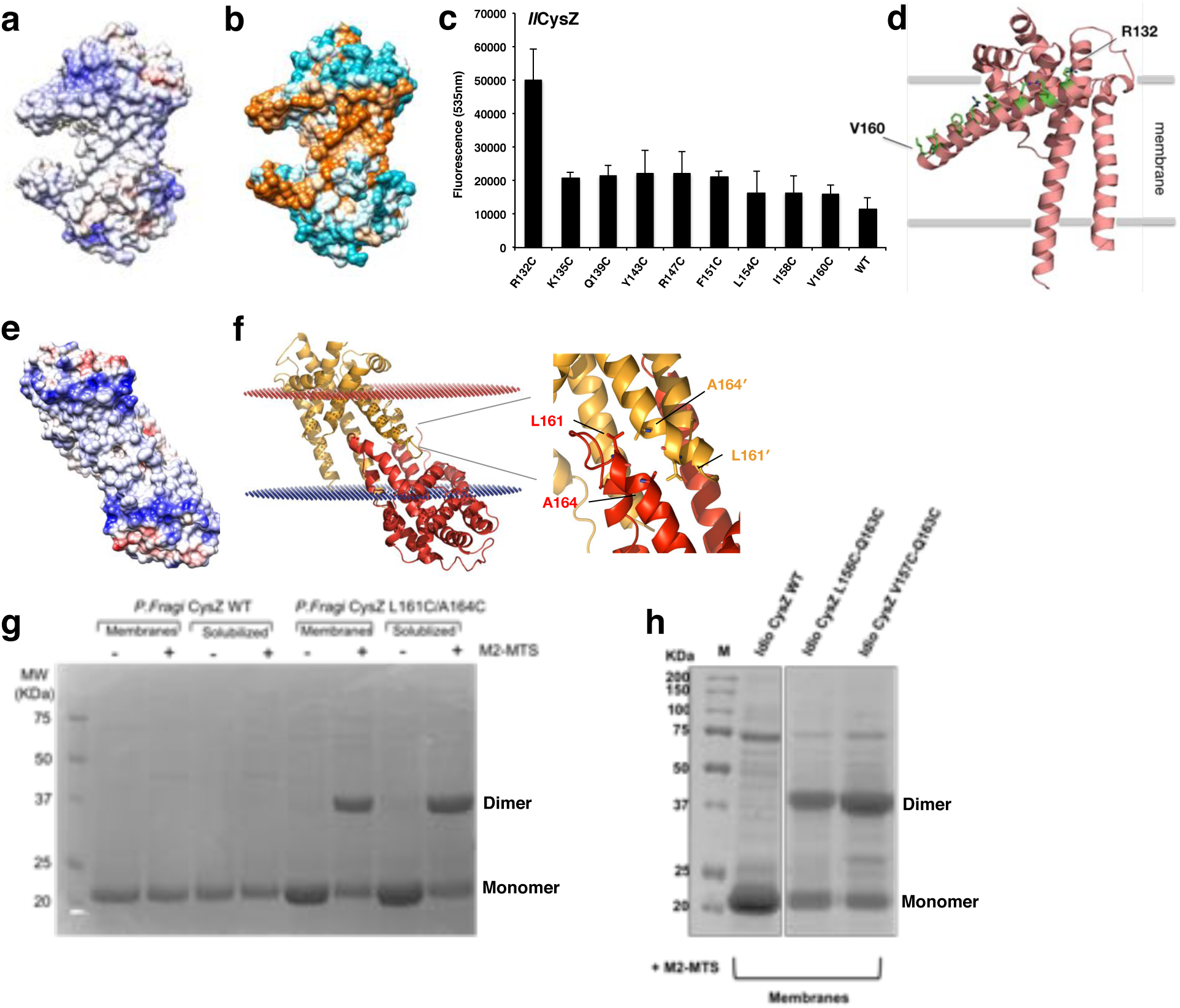
Surface electrostatics, hydrophobicity and site-directed fluorescence labeling of cysteine mutants located on H4 of *Il*CysZ. a. A depiction of the surface electrostatic potentials of the *Il*CysZ dimer with negative surface potential represented in red, and positive potential in blue as calculated by APBS. b. Surface of *Il*CysZ rendered by level of hydrophicity, using the Kyte-Doolitle hydrophobicity scale with cyan being most polar, to orange being most hydrophobic, reveals a hydrophobic belt along the center of the dimer, suggesting the orientation of the dimer in the lipid bilayer. c, d. Single cysteine mutants were designed to be located along the length of helix 4 of CysZ, to gauge the extent of its membrane insertion. The labeled protein was extracted from the membrane and purified after quenching the labeling reaction. Fluorescence intensity was measured and quantified by a Tecan fluorescence plate reader at an excitation of 485nm and emission of 535nm the results of which were plotted on the left, with error bars representing the standard deviation from the mean for n=3. The results show that of all the residues on the helix, only R132C (the top-most residue) was accessible to the fluorophore and hence exposed out of the membrane. d. Locations of the mutated residues on helix H4 are marked on the model of *IlCysZ* in green. Crosslinking of L161C-A164C cysteine mutant of *Pf*CysZ and *Il*CysZ exhibits dimer formation. e. A depiction of the surface electrostatic potentials of the *Pf*CysZ dimer with negative surface potential represented in red, and positive potential in blue as calculated by APBS, highlights hydrophobic belt marking the orientation of the *Pf*CysZ dimer in the membrane. f. Membrane orientation of *Pf*CysZ predicted by OPM/PPM server shows its 31-degree tilt to the perpendicular, with a zoomed in view of the location of the cysteine mutants at the dimer interface, used for crosslinking of the dimer. g. h. Dimer formation by the crosslinking of *Pf*CysZ (g) and *Il*CysZ (h) with introduced structure-based cysteine substitution mutations in the H4-H5 dimer interface. Analogous pairs of cysteine mutants from each species are shown here: *Pf*CysZ L161C-A164C and *Il*CysZ L156C-Q163C and V157C-Q163C. Sulfydryl specific Bis-MTS crosslinkers of a certain spacer length (in this case 5.2Å) were used at 0.5mM for 1hour at room temperature to crosslink the protein. Experiment was performed on protein in the membrane, as well as on solubilized protein. WT protein (cysteineless) was used as a control. Proteins were purified after stopping the reaction, and run on an SDS PAGE with reducing dye, and stained with Coomassie blue.

**Fig. 4. S1.**
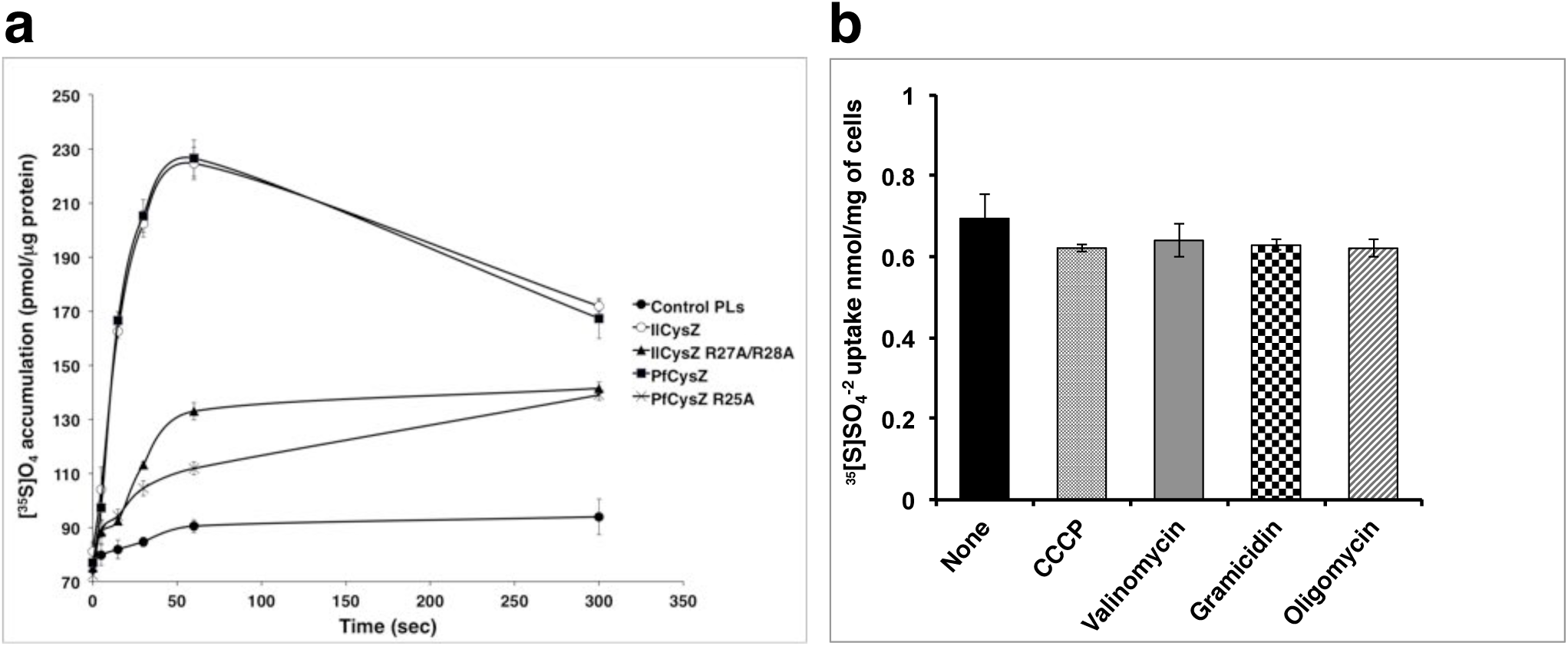
a. Radiolabeled sulfate (10 *μ*M [^35^S]O_4_ ^2-^) accumulation is measured from detergent purified *Il*CysZ and *Pf*CysZ reconstituted in proteoliposomes, in comparison to the control (empty) liposomes. Sulfate-binding site alanine mutants (*Il*CysZ R27A/R28A and *Pf*CysZ R25A) display a significantly diminished sulfate uptake capability. Error bars represent the standard deviation from the mean (SEM), for n=3. b. Effect of uncouplers and ionophores on the sulfate accumulation by CysZ. *Il*CysZ expressed in the *E. coli* knockout strain shows unchanged sulfate uptake levels in the presence of the uncouplers CCCP and oligomycin or the ionophores gramicidin and valinomycin. Sulfate accumulation was measured in triplicate at pH 7 and total accumulation was plotted after a 10 s incubation. Error bars were calculated by standard deviation from the mean, for n=3.

**Fig. 4. S2.**
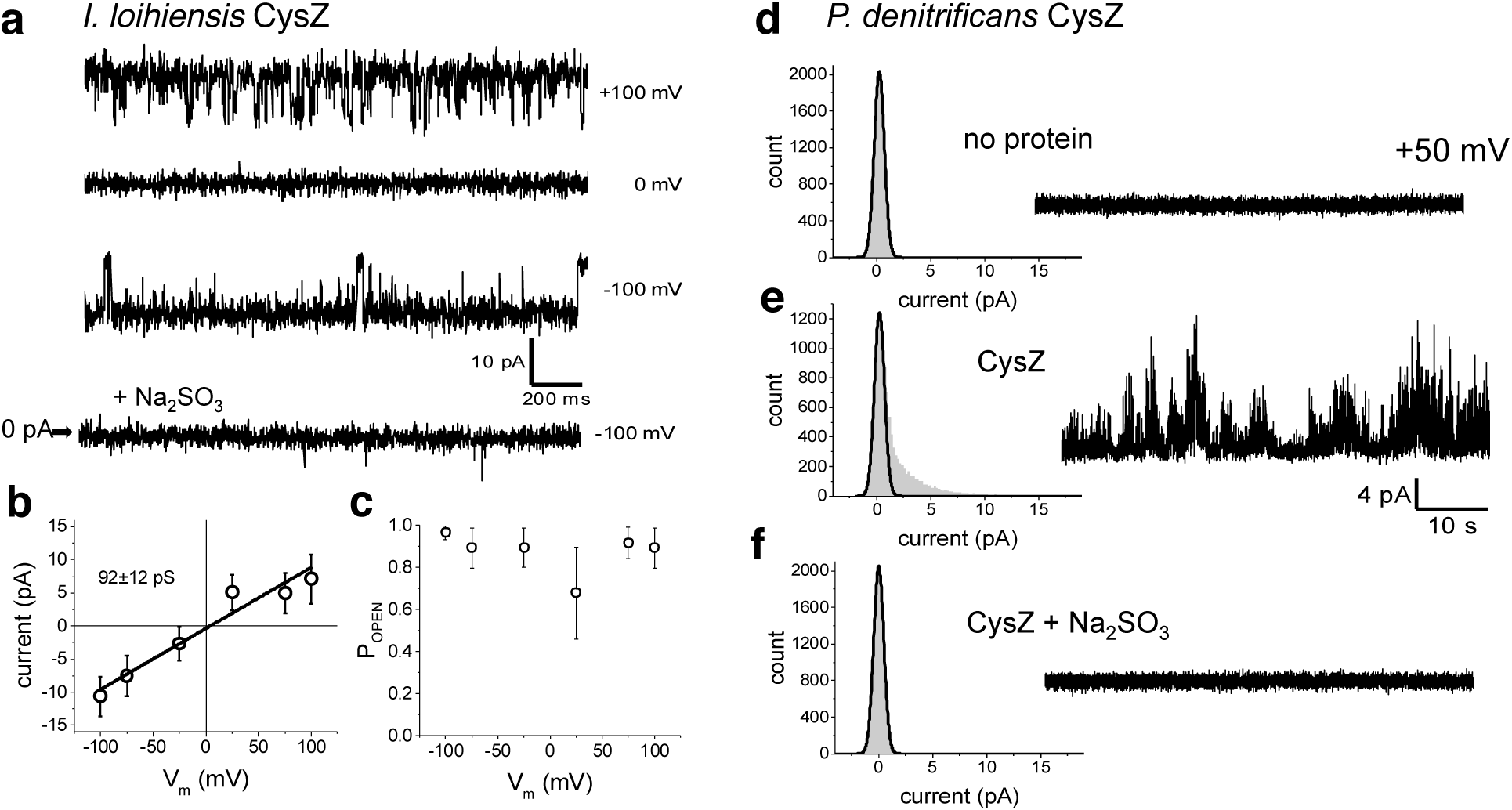
Reconstitution of CysZ in planar lipid bilayers yields sulfite-sensitive conductances. a. Sample traces of *Il*CysZ under 150 mM symmetrical Na_2_SO_4_ solutions at the indicated membrane potentials and pH = 5.4. The lowest trace shows inhibition of activity by 15 μM Na_2_SO_3_. b. Unitary conductance estimated at 92 ± 12 pS with a reversal potential (E_rev_) at 0 mV. c. Unitary open probability does not change as a function of membrane potential, and is stable at an average P_OPEN_= 0.8. d. Stable bilayer recording at a membrane potential V_m_ = +50 mV, with no protein reconstituted. An all point current amplitude histogram is plotted on the left of the trace. The peak of the normal distribution denotes the baseline zero current. e. Reconstitution of *Pd*CysZ under 150 mM symmetrical Na_2_SO_4_ solutions at +50 mV. An all point current amplitude histogram is plotted on the left of the trace. The high frequency activity shows a poorly resolved distribution of unitary currents above baseline. f. Inhibition of activity by 5 μM Na_2_SO_3_. An all point current amplitude histogram is plotted on the left of the trace. Sulfite inhibited activity back to baseline levels.

**Fig. 7. S1.**
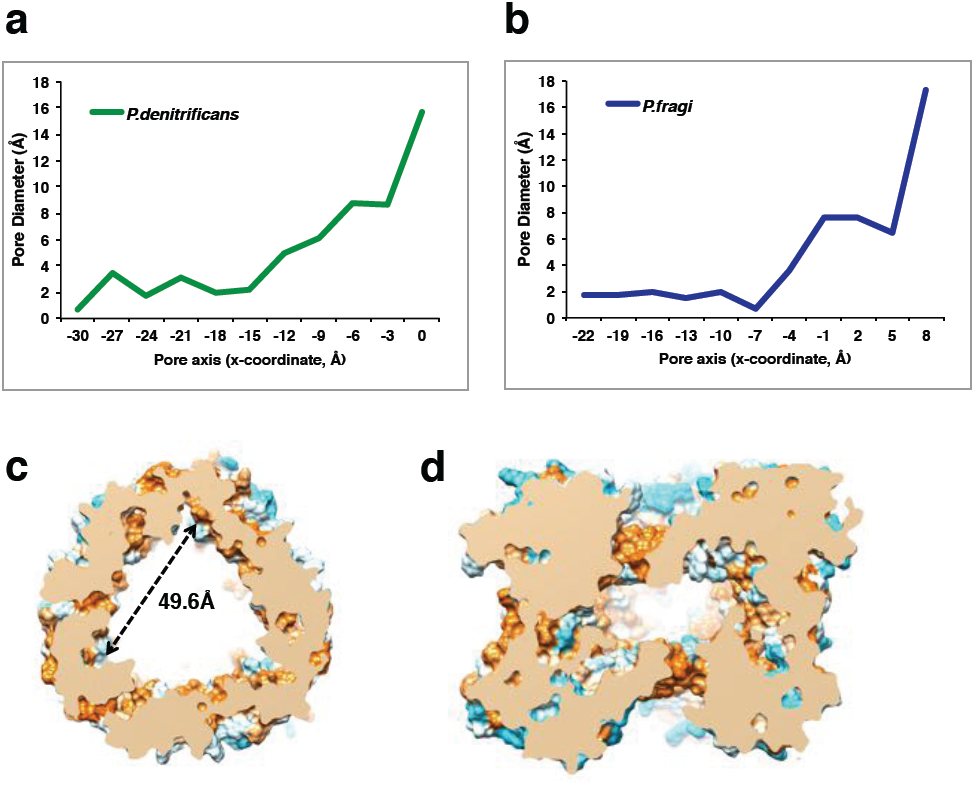
Plot of the pore diameter of the putative ion conduction pathway. a. *Pd*CysZ b. *Pf*CysZ, showing a very narrow constriction of under 2 Å in diameter near the entrance of the pore, which then widens as the ions move past the core of the protomer to the inner central cavity of the hexameric channel (as calculated by PoreWalker, measured at 3 Å steps along the x-axis) c. Central hydrophobic cavity of *Pd*CysZ hexamer, cross-section of central cavity of the hexameric *Pd*CysZ (top view) and d. side view, shows a triangular-shaped cavity with each side measuring 49.6 Å in length. The cavity is primarily hydrophobic, as depicted and colored with the Kyte-Doolitle hydrophobicity scale from most polar colored in cyan to most hydrophobic colored in orange.

